# The distribution of beneficial mutational effects between two sister yeast species poorly explains natural outcomes of vineyard adaptation

**DOI:** 10.1101/2024.06.03.597243

**Authors:** Emery R. Longan, Justin C. Fay

## Abstract

Domesticated strains of *Saccharomyces cerevisiae* have adapted to resist copper and sulfite, two chemical stressors commonly used in winemaking. *S. paradoxus*, has not adapted to these chemicals despite being consistently present in sympatry with *S. cerevisiae* in vineyards. This contrast represents a case of apparent evolutionary constraints favoring greater adaptive capacity in *S. cerevisiae*. In this study, we used a comparative mutagenesis approach to test whether *S. paradoxus* is mutationally constrained with respect to acquiring greater copper and sulfite resistance. For both species, we assayed the rate, effect size, and pleiotropic costs of resistance mutations and sequenced a subset of 150 mutants isolated from our screen. We found that the distributions of mutational effects displayed by the two species were very similar and poorly explained the natural pattern. We also found that chromosome VIII aneuploidy and loss of function mutations in *PMA1* confer copper resistance in both species, whereas loss of function mutations in *REG1* were only a viable route to copper resistance in *S. cerevisiae*. We also observed a single *de novo* duplication of the *CUP1* gene in *S. paradoxus* but none in *S. cerevisiae*. For sulfite, loss of function mutations in *RTS1* and *KSP1* confer resistance in both species, but mutations in *RTS1* have larger average effects in *S. paradoxus*. Our results show that even when the distributions of mutational effects are largely similar, species can differ in the adaptive paths available to them. They also demonstrate that assays of the distribution of mutational effects may lack predictive insight concerning adaptive outcomes.

## Introduction

Adaptation is limited by the variation mutations can provide. Historically, it has often been asserted that mutations can yield variation along any dimension of an organism’s phenotype (Lewontin 1974, Futuyma 1979, Futuyma 2010). This assumption in large part stems from the widespread success of artificial selection in laboratory environments and in domesticated plants and animals (Dobzhansky et al. 1977, Falconer and Mackay 1983, Hill and Caballero 1992). Consequently, influential models of adaptation such as Fisher’s geometric model have focused on pleiotropy as the major source of evolutionary constraints rather than mutational availability (Fisher 1930, Orr 1998, Orr 2005). However, failures of the adaptive process due to lack of variation are not as uncommon as might be expected under this framework. For example, many plant species in England have failed to adapt to heavy metal contamination near copper mines whereas others have succeeded (Bradshaw 1984, Bradshaw 1991). Additionally, artificial selection experiments on some traits such as desiccation resistance in *Drosophila birchii* fail to elicit any response (Hoffmann et al. 2003). There are also intrinsic physical constraints on the structures of organisms that preclude improvement by mutation beyond certain thresholds (Alexander 1985). Failure of adaptation due to lack of variation, termed “genostasis” or “sluggish evolution”, challenges the idea that mutations can always enable adaptive evolution (Bradshaw 1991, Futuyma 2010).

The distribution of mutational effects (DME) is the distribution of effects for all possible mutations an organism or species can experience weighted by their frequency. In principle, the DME itself defines constraints on adaptation for the first step along an adaptive walk (Couce et al. 2024). Theory predicts that the distribution of effect sizes for beneficial mutations should be exponential (Gillespie 1991, Orr 2003, Eyre-Walker and Keightley 2007), and there have been several empirical studies which attempted to assay the DME, sometimes referred to as the distribution of fitness effects (Wloch et al. 2001, Sanjuan et al. 2004, Böndel et al. 2019, Couce et al. 2024). In general, what is found is that most mutations are slightly deleterious or neutral, a large fraction of the remainder are lethal mutations, and a small fraction of mutations are beneficial to varying degrees. The small fraction of beneficial mutations is both critical to adaptation and very difficult to assay in most systems. This makes it hard to answer questions about the relative roles of chance versus adaptive potential when studying constraints in related species, particularly when one of the species adapts more successfully than another. As noted by Bradshaw (1991), “that what may occur in one organism, population, or species, may not occur in another”, but the role of mutational availability in determining these outcomes is only knowable if the beneficial tail of the DME is empirically measured.

Currently, there is a large gap in the understanding of how closely related species may differ in their adaptive capacity due to mutational constraints. This knowledge gap is particularly relevant to shifts in the environment that are universal to all resident species, such as changes in temperature or precipitation, or the introduction of a novel chemical stress. In such cases, there are many examples of successful parallel adaptation that span extreme phylogenetic distances, such as the evolution of cyclodiene resistance in four separate orders of insects by a parallel amino acid substitution (ffrench-Constant 1994). However, in cases like the metal contaminated soils mentioned above, only a subset of plant species were able to successfully adapt to the excess copper introduced to the soil (Bradshaw 1984, Bradshaw 1991). Examples like this illustrate that related species can differ drastically in their adaptive capacity and their evolutionary constraints. As of now, evolutionary biology has very little power to predict if adaptation in any particular species will fail or succeed in these types of scenarios (Futuyma 2010). Recent comparative experimental evolution studies have shown that related species often take similar, though not identical evolutionary paths towards adapting to a particular stress (Sanchez et al. 2017, Pentz and Lind 2021, Pentz et al. 2024). However, these studies are difficult to interpret from a mutational availability perspective because the outcome of experimental evolution is driven by interactions between mutation and selection (Van den Bergh et al. 2018). In the present study, we used mutagenesis as an alternative method to directly assay the mutational contribution to constraints.

As global change (Rey et al. 2016) and the mass extinction accompanying it progress, differential constraints will only become more important determinants of persistence and extinction (Cowie et al. 2022). A better understanding of mutational constraints is also needed regarding pathogens and their relatives because it remains difficult to predict which pathogens can most effectively adapt to particular treatments. Strikingly, related bacterial species have been shown to vary markedly concerning their capacity to adapt to certain drugs (Vogwill et al. 2014, Vogwill et al. 2016). Constraints also play key roles in the evolution of immune evasion for viruses such as influenza and SARS-CoV2, and the utility of predictive knowledge concerning these constraints cannot be understated (Wu and Wilson 2017, Carabelli et al. 2023).

We have taken a comparative mutagenesis-based approach to assay the DME with specific emphasis on beneficial mutations in two yeast species that differ in their adaptation in nature to two anthropogenic stressors relevant to winemaking. The goal of this work is to test the explanatory power of the DME in this natural case of apparent constraints and to determine how the DMEs of these two related species differ regarding two stressors that are important in the vineyard environment. *Saccharomyces cerevisiae*, unlike its sister species *S. paradoxus*, has successfully adapted to the enological stressors copper and sulfite. These species are routinely found in sympatry in vineyard environments and in oak forests (Sniegowski et al. 2002, Redžepović et al. 2002, Hyma and Fay 2013, Dashko et al. 2016, Vaudano et al. 2019). Domesticated strains of *S. cerevisiae* show much higher levels of copper and sulfite resistance compared to their wild counterparts (Pérez-Ortin et al. 2002, Fay et al. 2004, Warringer et al. 2011, Clowers et al. 2015, Dashko et al. 2016). *S. paradoxus* is widely considered a non-domesticated species, and despite being exposed to these chemicals in the vineyard environment, remains nearly universally sensitive to them (Liti et al. 2009, Warringer et al. 2011, Dashko et al. 2016, Yue et al. 2017).

Copper has been used as an agricultural antimicrobial since at least the mid 1700s (Money 2006, Borkow and Gabbay 2009) and came to prominence in viticulture in the 1880s when Pierre Marie Alexis Millardet published his experiments on the control of downy mildew and powdery mildew with a mixture of copper sulfate and calcium hydroxide which he called the “Bordeaux mixture” (Millardet 1885, Dixon 2004, Ayres 2004, Money 2006). Due to its low cost, high effectiveness, ease of use compared to other similar mixtures, and lack of adaptation in the problematic mildews it was intended to combat, this mixture has been sprayed in many vineyards yearly since the end of the 19th century (Masson 1887, Ayres 2004, Gessler et al. 2011). This has resulted in very high levels of copper in vineyard soils and alterations to microbial ecology in the affected areas (Besnard et al. 2001, Dell’Amico et al. 2008, Komárek et al. 2008, Mackie et al. 2012, Mackie et al. 2013, Fernández-Calviño and Bååth 2016, Grangeteau et al. 2017). The other primary use of copper in winemaking is the practice of adding copper sulfate to both red and white wines to remove sulfidic off-flavors, though this is not presumed to be a major source of selection on yeasts (Clark et al. 2015, Vela et al. 2017, Echave et al. 2021).

In the 1980s, it was discovered that high levels of copper tolerance in *S. cerevisiae*, a trait that is presumed to be uncommon in microbes (Borkow and Gabbay 2009), was due to tandem amplification of the *CUP1* gene which encodes a short, cysteine-rich metallothionein that sequesters copper ions in the cytoplasm (Fogel and Welch 1982, Fogel et al. 1983, Capdevilla et al. 2012). *CUP1* copy number correlates quite strongly, though not perfectly, with copper resistance in *S. cerevisiae* (Fogel and Welch 1982, Fogel et al. 1983, Warringer et al. 2011, Adamo et al. 2012, Gerstein et al. 2015, Strope et al. 2015, Hull et al. 2017). Sequestration of excess copper is critical because of the detrimental effects free copper can have including disruption of iron-sulfur clusters (Macomber and Imlay 2009), destabilizing the cell membrane via lipid peroxidation (Avery et al. 1996), and damaging DNA (Tkeshelashvili et al. 1991). Interestingly, tandem amplification of this gene has independently occurred at least five times in *S. cerevisiae* and has never been observed in *S. paradoxus* (Welch et al. 1983, Zhao et al. 2014, Yue et al. 2017). Although *CUP1* amplification is not the only genetic contributor to copper resistance in *S. cerevisiae* (Welch et al. 1989, Culotta et al. 1994, Gerstein et al. 2012, Chang et al. 2013), its absence in *S. paradoxus* likely explains a large fraction of the species difference.

Sulfite has been used as an antimicrobial in winemaking since at least the 19^th^ century (Divol et al. 2012). Claude Ladrey made mention of burning sulfur in barrels as early as 1871 in his book about winemaking (Ladrey 1871, Divol 2012). By the middle of the 20th century sulfite addition was common practice in the wine industry to suppress unwanted microbial growth during fermentation and prevent spoilage (Amerine et al. 1972, Jolly et al. 2006). There is evidence for the use of sulfur as an antimicrobial in winemaking long before this in ancient Egypt and in the Roman Empire, though the exact history is less clear than for copper (Pecci et al. 2020). Many domesticated strains of *S. cerevisiae* are very tolerant of sulfite due to high expression of the *SSU1* gene, which encodes a sulfite efflux pump. This high expression has evolved independently at least three times and, in all cases, involves structural variations that alter the upstream sequence of *SSU1* (Goto-Yamamoto et al. 1998, Pérez-Ortin et al. 2002, Zimmer et al. 2014, García-Ríos et al. 2019). Similarly, *S. uvarum*, a more distantly related *Saccharomyces* yeast, has also convergently evolved greater *SSU1* expression via rearrangements, but no known strain of *S. paradoxus* has done so (Macías et al. 2021). Of note, it has been shown recently that high expression of *SSU1* has an intrinsic trade-off with copper resistance due to Cup1 and Ssu1 having biochemically antagonistic roles in sulfur assimilation (Onetto et al. 2023). However, many domesticated strains retain resistance to both chemicals well beyond that of their wild counterparts (Dashko et al. 2016).

Sulfite tolerance is quite rare among other microbes (for some exceptions see Stratford et al. 1987, Varela et al. 2019). Although there is some variation within *S. paradoxus*, studies that compare the two species consistently find that *S. cerevisiae* is far more sulfite resistant than the near-universally sensitive species *S. paradoxus* (Dashko et al. 2016). Given that *S. paradoxus* is routinely isolated from vineyards and wine must, these phenotypic differences suggest that *S. paradoxus* may be mutationally constrained with respect to its adaptation to sulfite.

Using this case of differential adaptation between microbial species as a model, we took a mutagenesis-based approach to assay the DME between species to determine if differences in the DME may have predisposed the outcome seen in nature, namely *S. cerevisiae* repeatedly adapting and *S. paradoxus* repeatedly failing to adapt. If copper and sulfite resistance mutations are (1) more abundant, (2) of larger effect, or (3) tend to come with fewer pleiotropic costs in *S. cerevisiae* than in *S. paradoxus*, then the DME could explain why we see successful adaptation in the former species but not the latter. Many studies have used microbial experimental evolution to ask similar questions about species differences in the DME and its impact on adaptation. These studies have revealed that genetic background can play large roles in the frequency (Blount et al. 2008), effect size (Vogwill et al. 2014, Gonzalez and Bell 2013, Sanchez et al. 2017), and pleiotropic costs (Gagneux et al. 2006, Vogwill et al. 2016) of beneficial mutations. Such differences in the DME among genetic backgrounds have been termed “macroscopic epistasis” and represent a major avenue by which differences in the DME can lead to different evolutionary responses, and in extreme cases, determine the success or failure of adaptation (Good and Desai 2015).

In contrast to experimental evolution, mutagenesis-based techniques offer the unique advantage of circumventing the waiting time for adaptive variants to arise and fix. This advantage is exacerbated when comparing mutagenesis to experimental evolution of large asexual populations where clonal interference can greatly slow the rate of adaptation and fixation events (Gerrish and Lenski 1998, de Visser and Rozen 2006, Perfeito et al. 2007, Maddamsetti et al. 2015). Additionally, mutagenesis-based techniques allow for rare, large effect mutations to be sampled more completely than in experimental evolution (Wloch et al. 2001). In this study, we UV mutagenized copper and sulfite sensitive strains of *S. cerevisiae* and *S. paradoxus* and directly recovered mutants from plates containing various concentrations of these chemicals to assay mutational target size. We then subjected thousands of these mutants to a high-throughput phenotyping assay to measure mutational effect size and pleiotropic costs. A smaller subset (N = 150) had their genomes sequenced, and we identified causal variants underlying resistance in both species.

Contrary to expectations, we find that differences in mutational target size and mutational effect size do not align with the pattern seen in nature. Also, patterns of pleiotropic costs only show a slight trend favoring greater costs in *S. paradoxus* copper mutants. In the sequenced strains we find that the same genes (*PMA1* for copper and *KSP1* and *RTS1* for sulfite) and chromosome aneuploidy (Chr VIII) tended to be mutational targets in both species. However, there are many important differences between the two DMEs in terms of the mutations recovered for each species, along with their individual effect sizes and costs. Overall, the results of this study corroborate the notion that mutational effects tend to be generally but not universally conserved between related species. These results also highlight that the DME on its own may sometimes offer very little predictive or explanatory power when considering cases of apparent constraint, suggesting that evolutionary potential cannot be reliably inferred from measures of the DME.

## Materials and Methods

### Yeast strains

We generated the focal strains used in this study from two *S. cerevisiae* and two *S. paradoxus* strains isolated from trees in Greece and France respectively (Robinson et al. 2016, Table S1). The two *S. cerevisiae* strains are members of a Mediterranean oak population from which domesticated wine strains are thought to have been derived (Almeida et al. 2015). We knocked out the *HO* gene in monosporic derivatives of each strain by transforming a natMX4 deletion cassette constructed using the plasmid pAG25 and primers that targeted the *HO* locus (Goldstein and McCusker 1999, Table S2). For the *S. cerevisiae* strains, we used primers JM7 and JM8 from Goldstein and McCusker (1999), and for *S. paradoxus* we designed analogous primers with homology to the *S. paradoxus HO* locus (Table S2). All transformations in this study were carried out using the lithium acetate/salmon sperm/PEG method (Geitz and Schiestl 2007). For all transformations into *S. paradoxus*, a modified heat shock of 40°C was used. We next sporulated transformants and dissected tetrads to obtain haploid derivatives of our ancestral strains. Successful *HO* integration was confirmed via co-segregation of nourseothricin resistance and haploidy assayed via PCR amplification of the *MAT* locus (Huxley et al. 1990, Table S2).

For each monosporic clone, *MAT*a and *MAT*α haploid strains were obtained (Table S1). These haploids were then mated to their isogenic counterparts on YPD (Bacto yeast extract 10 g/L, Bacto peptone 20 g/L, dextrose 20 g/L, agar 20 g/L) plates to obtain homozygous diploids, which were confirmed as diploid via PCR amplification of the *MAT* locus and the capacity to sporulate. Following these manipulations, our focal strain set for this study included 12 strains derived from the four wild isolates: A *MAT*a haploid derivative, a *MAT*α haploid derivative, and a homozygous diploid derivative of each (Table S1). In our phenotyping assays, derivatives of the laboratory *S. cerevisiae* strain S288c (YJF5538) and a North American oak *S. cerevisiae* strain, YPS163 (YJF5539), were included as points of reference (Mortimer and Johnston 1986, Sniegowski et al. 2002).

In addition to the focal strains, 150 mutants derived from these strains were sequenced and are listed in Table S3. For follow up assays on *PMA1* mutations, five heterozygous diploid strains for each species were constructed by mating sequenced mutants to their *MAT*α parent (Table S1). For follow up assays on *REG1*, deletion strains of both species were constructed via transformation of an hphMX4 cassette targeted to the *REG1* locus (Table S2). Deletion mutants were confirmed via PCR. A suppressor mutant of the slow growth phenotype seen in the *S. paradoxus REG1* deletion strain was also recovered and included in the *REG1* follow up assays (Table S1). Information concerning the other mutants used in the high throughput phenotyping and the data relevant to them can be found with the following DOI: 10.6084/m9.figshare.25777512 .

### Mutagenesis

We mutagenized two haploid (*MAT*a) and two diploid strains from each species. Our mutagenesis protocol is similar to the irradiation protocol found in Birrel et al. (2001). We used 2 mL of saturated overnight culture (YPD) as our input into mutagenesis and our control mock mutagenesis for each strain. This culture was spun down at 700 rcf for five minutes and was resuspended in 10 mL of autoclaved deionized water. This suspension was then transferred to a petri dish and subjected to either 0 seconds (mock mutagenesis) or 10 seconds of UV-C radiation in a Singer Rotor HDA (Singer Instruments, Somerset, England). The cell suspension was then moved back to a 15 mL conical tube and spun down at 700 rcf for five minutes. Next, we resuspended the mutagenized and control pools into 6 mL of YPD and allowed the cultures to recover overnight at 30°C with shaking at 250 rpm and then stored these cells the following day as 15% glycerol stocks at −80°C.

### Mutant isolation

From the stocks of mutagenized and mock mutagenized cells, we measured the induced mutation rate and recovered mutants with elevated copper and sulfite resistance. To accomplish the former, we used a canavanine plating assay. Canavanine is an arginine analog that is toxic to wild type cells. However, loss of function mutations in the arginine transporter *CAN1* confer resistance to the drug. This allows canavanine to be used as a control for induced mutation rate in haploids. To isolate copper and sulfite mutants, we plated the mutagenized and mock mutagenized pools onto solid media containing added copper or sulfite. Specifically, 200 μl of the mutagenized and control pools for both the haploid and diploid strains were plated on eight concentrations of complete media (CM: US Biological drop-out mix complete without yeast nitrogen base 1.3 g/L, Difco yeast nitrogen base 1.7 g/L, 5 g/L ammonium sulfate, dextrose 20 g/L, agar 20 g/L) + copper sulfate (0.0 mM, 0.05 mM, 0.1 mM, 0.15 mM, 0.2 mM, 0.3 mM, 0.4 mM, 0.5 mM, and 0.6 mM). Additionally, we plated the pools on eight concentrations of CM + 75 mM tartaric acid/sodium tartrate (pH = 3.5) with various amounts of added 0.5 M sodium sulfite. The final concentrations were: 0 mM, 0.5 mM, 1 mM, 1.5 mM, 2 mM, 3 mM, 4 mM, 5 mM, and 6 mM sulfite. Notably, sodium sulfite has long been known to lack thermostability due to off-gassing of the main cytotoxic sulfite species, SO_2_, complicating formation of solid sodium sulfite plates. Our method was to add sodium sulfite to 50 mL aliquots of hot autoclaved agar media and to pour the plates immediately. This consistently led to toxicity of sulfite being retained and to apparent uniformity of sulfite concentrations within single plates as compared to the potentially noisier methods that have been used in the past such as spreading sodium sulfite onto solidified plates (Park et al. 1999). To control for cell density, 200 μl of a 10^-5^ dilution of each mutagenized and mock mutagenized haploid pool were plated onto YPD. Plating 200 μL of a 10^-4^ dilution was used for this purpose for diploids. After plating, plates were incubated at 30°C for five days and the number of colonies per plate was counted (Table S4).

### Mutation rate calculation

We estimated the induced mutation rate in our haploid mutagenized pools by plating mutagenized and control pools of each haploid strain on three concentrations of CM + canavanine (30, 45, and 60 mg/L). The mutant induction rate was calculated as the number of colonies recovered divided by the total number of cells plated for each strain on each concentration of canavanine. To assay differences among strains, we compared our canavanine mutation rates across the three concentrations between species with a paired t-test. We also compared mutation rates at the highest concentration with a chi-square test. Next, using canavanine mutation rate estimates from Lang and Murray (2008), we estimated the mutations per genome in our mutagenized pool along with the saturation of our screen (see also Gruber et al. 2012, Metzger et al. 2016, and Hodgins-Davis et al. 2019 for similar calculations). Using a canavanine plating assay, Lang and Murray (2008) estimated the number of spontaneous mutations per base pair per cell division as 6.44 x 10^-10^ based on the rate at which they recovered mutants with a canavanine resistance phenotype, which was 1.52 x 10^-7^. We used these rates to estimate the number of induced mutations per genome by multiplying this per base pair mutation rate by the fold increase in phenotypic mutation rate we observe in our mutagenized pools on 60 mg/ml canavanine relative to Lang and Murray’s phenotypic mutation rate (Table S4). Across the four strains we saw an average of an 88-fold increase in phenotypic mutation rates yielding an estimate of 5.66 x 10^-8^ mutations per base pair. When multiplied by the length of the genome this yields an average of 0.68 mutations per genome in our mutagenized pools. When our inferred per base pair mutation rate is multiplied by the number of cells plated in each screen it yields an average of 1.7 mutations per base pair in the genome for copper and 1.22 mutations per base pair in the genome for sulfite. These numbers refer to the average number of mutants in the whole screen at any given site in the yeast genome (Table S4). Critically, these numbers do not take into consideration the biased mutation spectrum of UV mutagenesis (Mao et al. 2017). For our plating assays on copper and sulfite, mutation rates were simply calculated as the number of colonies recovered divided by the total number of cells plated for each strain on each concentration. These rates were compared strain to strain using paired t-tests and chi-square tests where appropriate. Colony counts and inferred saturation calculations can be found in Table S4.

### Mutant picking

A subset of haploid (N = 3,024) and diploid (N = 720) mutants that produced colonies on stress concentrations that killed the ancestor were sampled for follow up phenotyping to assess the effect size and pleiotropic consequences of the mutations. Sampling was done by recovering single colonies via pipette tips and manually arraying cells from these colonies in a grid on a solid CM plate. For haploid copper mutants, we sampled 1,512 resistant isolates and arrayed them in four (one for each ancestral strain) 384 format plates along with 6 controls per plate. These controls were YJF5538 (S288c derivative), YJF5539 (YPS163 derivative), two replicates of the ancestral strain, and two independent isolates recovered from the mutagenized pool without selection to control for the effect of mutagenesis. The mutants consisted of 168 spontaneous mutants and 1,344 induced mutants. For haploid sulfite mutants, the same procedure and controls were used to array 1,025 spontaneous mutants and 487 induced mutants. Regarding sulfite mutants, both *S. paradoxus* strains produced fewer than 378 mutants (N = 330 for YJF3734 and N = 110 for YJF3815). Thus, the remaining positions on these plates were filled with *S. cerevisiae* derived sulfite mutants. After initial arraying on 384 plates, these were collapsed to 1536 format solid plates to await phenotyping. Figure S1A summarizes the mutant picking pipeline.

For diploid copper and sulfite mutants, we similarly sampled 90 resistant isolates for each of the strains from the plating assay. These were arrayed in 96 format along with six controls, which were four replicates of the ancestor, and a replicate each of YJF5538 and YJF5539. For both copper and sulfite, these mutants comprised 48 spontaneous mutants and 312 induced mutants split evenly among the four ancestral strains. These plates were collapsed to 1536 format with two technical replicates of each mutant on the plate to await phenotyping. For both diploids and haploids, mutants appearing on higher concentrations of copper and sulfite were prioritized and sampled exhaustively because these were putatively the largest effect mutants. We sampled as evenly as our plates permitted across the lower concentrations. For each mutant, we recorded three aspects of colony morphology when picking them from the stressor plate: relative size, color, and circularity. These data can be found at DOI : 10.6084/m9.figshare.25777512 .

### Phenotyping

Arrayed strains were phenotyped on copper, sulfite, and six permissive conditions. For both haploid copper and sulfite mutants, a master 1536 plate (CM) harboring all of the mutants and controls was replica plated onto phenotyping plates. Replica plating was performed using a Singer Rotor HDA robot (Singer Instruments). For copper we used 18 concentrations ranging from 0 mM to 0.8 mM and for sulfite we used 16 concentrations ranging from 0 mM to 6 mM for haploids and 21 concentrations ranging from 0 mM to 6 mM for diploids. Our permissive conditions encompassed three different base media: YP, CM, and minimal media (MM: 1.7 g/L yeast nitrogen base, 5 g/L ammonium sulfate) with either dextrose (2%) or glycerol (3%) as a carbon source. All stressor and permissive conditions were incubated at 30°C.

Colony size was measured using images obtained in a Singer Phenobooth using the Phenosuite software package (Singer Instruments). Colony size measurements were performed at 24 hour intervals for three days for copper and permissive conditions. For sulfite we imaged the plates for five days because a growth delay is a common phenotype associated with sulfite stress. Colony sizes are recorded as the number of pixels the colony occupies in an image. Example raw and processed images are shown in Figure S2. For haploids, images were taken at 1280 x 960 resolution and for diploids 4128 x 3096 resolution was used.

After imaging, we had obtained a total of 522,240 colony size measurements in the haploid dataset and 299,520 in the diploid dataset. Colony size measurements were subjected to manual quality control, and technical replicates in the diploid dataset were averaged to a single measurement. During manual curation, we removed 63,382 and 9,784 colony size measurements from the haploid and diploid datasets respectively. These removals were due to failure of a colony being transferred to a plate or excessive drying of an area of a plate. For sequenced mutants, copper and sulfite phenotypes were confirmed via repeating the phenotyping protocol in triplicate in 384 format. All three phenotyping datasets for the haploid, diploid, and sequenced mutants can be found at DOI : 10.6084/m9.figshare.25777512 .

### Colony size analysis

For copper and sulfite, we measured resistance by the area under the curve (AUC) as a function of concentration. AUC was estimated using the trapezoid rule implemented in the R package Pracma (Borchers 2019). Each colony size measurement on a stressor plate was normalized to the relevant no-stress condition for each mutant. The AUC phenotyping method is summarized in Figure S1B. Comparisons in the AUC metric were made using ΔAUC relative to the ancestral AUC via Kruskal-Wallis tests. For our triplicate assays on the sequenced mutants, we used the same method and averaged the technical replicates. Permissive phenotypes were quantified as growth relative to the ancestor. Thus, a score of 1 means that a strain produced a colony of equivalent size as compared to its ancestor. Edge effects (Baryshnikova et al. 2010, Zackrisson et al. 2016) were removed from the permissive growth data by a “layer” based normalization (Miller et al. 2022). This normalization entails grouping colonies by the number of positions that separate them from the edge of the plate. On a 1536 plate, this yields 16 distinct layers. Row and column normalization was not possible because row is confounded with species in these experiments.

Within our copper and sulfite data, several “mutants” appeared to phenocopy their ancestor in stress conditions. This indicated that these strains produced colonies in our plating assay for reasons other than harboring a mutation conferring heritable stress resistance. We categorized these strains as “physiological escapees” and defined them statistically as any mutant failing to yield an AUC not exceeding three standard deviations greater than that of the ancestral mean. These “escapees” were removed from all analyses of our assays. For haploid sulfite mutants, we were only able to rule out our sequenced mutants as non-escapees (see below), and these strains’ triplicate phenotypes were the only ones retained in those colony size analyses.

### Genome sequencing

For copper mutants, we selected 120 mutants for sequencing. These 120 were evenly distributed across our four haploid ancestral strains. For mutants derived from each ancestor, we divided mutants into quartiles based on their copper resistance (ΔAUC). The bottom quartile was removed from consideration, and 10 mutants were randomly selected from the top three quartiles for each ancestral strain’s haploid derivatives. For sulfite mutants, the top 30 AUCs for each haploid ancestral strain’s derivatives were chosen as sequencing candidates. After triplicate phenotyping, 31 strains remained as non-escapees, 12 from *S. cerevisiae* and 19 from *S. paradoxus*.

Following selection, genome sequencing was completed for these 151 haploid mutant strains and each of the four ancestral strains. One copper mutant had very low coverage and was eliminated from further analysis. For each of the strains that were subjected to whole genome sequencing, we extracted genomic DNA using a Yeastar genomic DNA kit (Zymo Research), prepared our libraries using a Nextera DNA Flex Library Preparation Kit (Illumina), and sequenced them as multiplexed libraries on a Hiseq X platform (150bp, paired end) via Novogene.

We mapped reads for each strain to the *S. cerevisiae* or *S. paradoxus* reference genomes (S288C_reference_sequence_R64-2-1 and *S. paradoxus* ultrascaffolds retrieved from the *Saccharomyces sensu stricto* database) using version 0.7.17 of the Burrows-Wheeler aligner (Scannell et al. 2011, Li 2013). Reads were marked duplicate with Picard tools version 2.12.0 and variants were called using version 4.1.7.0 of GATK using the HaplotypeCaller command. We applied six filters to our dataset: 1) Sites that had four or more unique genotype calls across all sequenced strains were removed due to low confidence. 2) Because all sequenced strains are haploid, sites lacking two different homozygous genotype calls were removed. 3) Sites where the ancestral strain had a heterozygous call were removed. 4) If the ancestral genotype call disagreed with all other calls at the site, that site was removed. 5) If >10% of the derivatives of any ancestor lack a call at a site, that site was removed only for that ancestral strain and its derivatives. 6) If a single site had more than one insertion/deletion (indel) called in multiple strains or had a genotype quality less than 10, it was removed due to low confidence. After applying these filters for the *S. cerevisiae* mutants, 700 sites were retained across the derivatives of both ancestral strains out of 91,840. After applying these filters for the *S. paradoxus* mutants, 334 sites were retained across the derivatives of both ancestral strains out of 33,723. Details can be found in Table S5.

Mutations were annotated for their functional effects on protein sequence using SNPeff (Version 4.3t, build 2017-11-24 10:18) along with genome annotation files downloaded from the *Saccahromyces* Genome Database (S288C_reference_sequence_R64-2-1) and the *Saccharomyces Sensu Stricto* Database (Scannell et al. 2011). Of note, *PMA1* and *PMA2* are erroneously swapped in the annotations in the *Saccharomyces sensu stricto* database for *S. paradoxus*. We corrected this issue for our identification of mutants.

Structural variation was assessed using the program Delly (version 0.8.7). We applied a similar set of filters to our structural variant calls as our SNPs and indels. Additional filters for low quality and imprecise calls were added, and we altered the requirement for two different homozygous calls to exclude duplications, as these may be expected to be called heterozygous. To be specific, this alteration involved removing any calls that were not singletons, and removing any heterozygous singletons that were not duplications. We also manually inspected the remaining variants for anomalous patterns such as the majority of stains being called heterozygous for a variant. Of a total of 51,804 variants, only 5 were retained after these filters were applied (Table S5). To ensure that *CUP1* tandem duplications were not missed, we manually inspected coverage data at the *CUP1* locus for all copper mutants. For the strain YJF4464, which harbors a *CUP1* duplication, we also mapped reads to a modified reference sequence containing an inverted repeat of the *CUP1* region to determine the structure of the repeat. The *SSU1* locus was also manually inspected in all sequenced sulfite mutants, and the *FZF1* locus was analyzed and inspected accounting for the introgressed *S. paradoxus* allele present in YJF3731 and its derivatives. After *PMA1* was identified as causal for copper, manual inspection of this gene yielded discovery of one additional mutation that was previously missed in YJF4433 and two synonymous mutations that were missed in YJF4439. These *post hoc* findings are noted in Table S3.

Karyotypes for each strain were obtained systematically via sequencing depth. Specifically, for each strain, depth was calculated for each 1kb window across the genome, and these numbers were averaged for each chromosome. Then the depth for the individual chromosomes was normalized to the coverage of the least covered chromosome. Relative read depth was then assessed and an output for the chromosome count for each of the 16 chromosomes was obtained assuming there is a single copy of the least covered chromosome. Relative read depth was calculated disregarding the rDNA cluster on chromosome XII. These karyotypes were then manually inspected and confirmed via assessment of the genome wide coverage. To account for the possibility of diploidization (Mable and Otto 2001, Zeyl 2003, Gerstein et al. 2006, Selmecki et al. 2015), separate estimates of chromosome number were obtained assuming a copy number of two for the least covered chromosome. A strain was considered as having evidence for diploidization if more than one aneuploid chromosome showed coverage values consistent with the diploid estimates. Nine strains had a coverage pattern consistent with diploidization and this is noted in the karyotype calls (Table S3). Three strains had anomalous coverage patterns not easily accounted for by diploidization and these are also noted in Table S3. Manual inspection did not reveal any cases of partial chromosome loss and revealed only one clear case of partial chromosome gain in *S. paradoxus* strain YJF4480 (Table S3).

### Simulations to assess gene/aneuploidy significance

To determine which genes were significant hits in our mutant screen, we performed *in silico* simulations of our experiment using the number of mutations detected in our dataset that alter protein sequences. Specifically, we used our empirical number of disruptive mutations (nonsynonymous, frameshift, nonsense, in-frame indels, and stop lost mutations) as input for these simulations along with the length of every gene in the genome for each species. The simulation “rains down” a number of mutations equal to the number observed in our experiments onto the coding sequences of the genome and counts the number of hits found in each gene. By performing this simulation 1,000,000 times per species we arrived at a null distribution for the number of mutations expected for each gene accounting for gene length. This in turn yields an empirical p-value for each gene as the fraction of simulations with equal or greater hits than the number of hits in the dataset. We assessed the significance of each gene by applying a Bonferroni correction (N = 6,696 for *S. cerevisiae* and N = 5,963 for *S. paradoxus*) to these empirical p-values. For both species, information for each gene with multiple hits in our experiment and its significance can be found in Table S6.

A similar procedure was carried out for chromosome aneuploidies. We used the number of aneuploid strains and the number of aneuploid chromosomes in each of these strains as input. For each iteration of the simulation, this number of strains with the specified number of aneuploid chromosomes had chromosomes randomly selected as aneuploid. To illustrate, for *S. cerevisiae* there were eight total aneuploid strains. Four of these strains had one aneuploid chromosome, and the other four had two, three, and seven aneuploid chromosomes respectively. For each iteration of the simulation, eight strains with these numbers of aneuploidies had their aneuploid chromosomes selected at random, and these simulated data were used to give an empirical p-value for each chromosome. To test for enrichment of specific chromosome combinations, namely CHRIII and CHRVIII, we simulated the number of aneuploidies observed in our dataset for each chromosome and counted the number of strains harboring both aneuploidies for each iteration. These empirical p-values were used to assess significance. Incidence data concerning all significant genes and aneuploidies are summarized in Figure S3. Statistics comparing incidence, effect sizes, and costs between species for significant genes and aneuploidies are summarized in Table S7.

### PMA1 pH sensitivity and dominance assay

To test the effects of *PMA1* mutations on low pH sensitivity, we spot diluted 39 *PMA1* mutants onto MM and low pH MM (MM + 10 mg/mL unbuffered tartaric acid, pH = 2.5). Cells from an overnight culture were serially diluted on each medium. Mutants were qualitatively scored based on their relative growth on the two media relative to the ancestor as either low pH sensitive or not low pH sensitive (Table S8). Another follow up was carried out to determine if *PMA1* mutations are dominant or recessive with respect to their effect on copper resistance and acid sensitivity. For five of the six cases cases where *S. cerevisiae* and *S. paradoxus* yielded site level parallel changes in *PMA1*, the haploid mutant was crossed to its *MAT*α parent strain yielding a diploid heterozygous *PMA1* mutant (Table S1). These strains were then spot diluted as described above onto MM, low pH MM, CM, CM + 0.02 mM CuSO_4_, CM + 0.05 mM CuSO_4_, CM + 0.1 mM CuSO_4_, and CM + 0.2 mM CuSO_4_. The heterozygotes were scored qualitatively relative to their ancestor and their haploid counterparts (Table S9). An additional summary of all the amino acid changes in *PMA1* we recovered along with their known phenotypes (Young et al. 2023) in prior studies can be found in Table S10.

### REG1 deletion and spot dilutions

Following sequencing, *REG1* was chosen for follow up experiments. *REG1* was deleted from one of the *S. cerevisiae* (YJF3732) and one of the *S. paradoxus* (YJF3734) strains. For *S. paradoxus*, two strains were recovered from this process. Firstly, we recovered a *REG1* deletion transformant with an extreme slow growth phenotype. Secondly, we recovered a derivative of this same transformant that had acquired a suppressor mutation that restored near wild type levels of growth on CM. To test for the phenotypic effects of *REG1* deletion in *S. cerevisiae* and *S. paradoxus*, we performed spot dilution assays on four separate media for these three deletion strains and their two ancestors. The media used were CM, CM + 0.1 mM CuSO_4_, MM, and low pH MM. Cells were spot diluted on each medium as described above and imaged using a Phenobooth following three days of growth (Singer Instruments). YJF5535 (*S. paradoxus* Δ*reg1*) also had a 10x concentrate of its overnight culture included on the plates due to its poor growth rate. Additionally, several sequenced *REG1* mutants (YJF4444, YJF4445, and YJF4452) were also assayed for their sensitivity to low pH media as described above.

## Results

### Mutant Screen

To test whether differences in adaptation to vineyard stressors between *S. cerevisiae* and *S. paradoxus* can be explained by differences in their DME, we UV mutagenized haploid and diploid derivatives of two Mediterranean oak strains of both species and isolated mutants with elevated copper and sulfite resistance. If *S. paradoxus* is more mutationally constrained with respect to adaptation to copper and sulfite, then at least one of the following must be true: (1) *S. paradoxus* has fewer beneficial mutations conferring resistance to these stressors available in its genome than *S. cerevisiae* (target size hypothesis), (2) mutations conferring elevated resistance to these stressors in *S. paradoxus* are of a smaller average effect size than in *S. cerevisiae* (effect size hypothesis), (3) mutations conferring resistance in *S. paradoxus* are more costly in permissive conditions than mutations conferring resistance in *S. cerevisiae* (costs hypothesis). To test the target size hypothesis, we screened our mutagenized pools for resistance mutations by plating them onto several concentrations of copper and sulfite. We also plated our haploid mutagenized pools onto three concentrations of canavanine to calculate an induced mutation rate and the per base pair genome wide saturation of our screen (see Methods).

In our canavanine control, we find that induced phenotypic mutations rates were consistent between species across our three assayed concentrations (Figure S4). There is no species difference among our haploid mutagenized pools in induced mutation rate across the three concentrations (paired t-test, p = 0.75) or when only the highest canavanine concentration is considered (chi-square test, p = 0.85). This is consistent with equal induction of mutations between the two species. Spontaneous rates of canavanine resistance across the three concentrations also did not vary with species (paired t-test, p = 0.61).

We used data from our highest canavanine concentration to estimate the genome-wide saturation of our mutant screen. Lang and Murray (2008) measured per base pair mutation rates in *S. cerevisiae* using canavanine resistance. These figures have been used by many other studies to infer per base pair saturation of a mutant screen (Gruber et al. 2012, Metzger et al. 2016, Hodgins-Davis et al. 2019). Using a similar methodology, we estimate that UV mutagenesis increased the mutation rate 88-fold and resulted in ∼0.68 mutations per genome. Given the number of cells per ancestral strain screened on copper (average of 3.1 x 10^7^) and sulfite (average of 2.3 x 10^7^), we estimate our screen included >1 mutation per site per genome for each ancestral strain. Although the UV mutation spectrum is biased for certain types of mutations (Mao et al. 2017), our calculations give rise to an expectation that each gene was mutated many times and site level parallelism had a reasonable probability of occurring (Table S4). *Mutational target size*

To estimate rates of copper and sulfite resistance we plated mutagenized and mock mutagenized cells on plates with a range of copper and sulfite concentrations. There were no differences between species in the recovery rates of induced or spontaneous mutants for either haploids or diploids on both stressors (Figure 1). On copper we obtained 8,704 mutants, mostly from the lowest concentrations. Both ancestral *S. cerevisiae* strains produced mutants that were recovered on higher concentrations than any of the *S. paradoxus* mutants. Also, for several of the lower concentrations, many more *S. cerevisiae* mutants were recovered than *S. paradoxus* mutants. However, when all concentrations were analyzed together, the frequencies of both induced and spontaneous mutants did not differ between species for either haploids or diploids (concentration paired t-test, p > 0.05 in all cases). The distribution of mutants recovered across concentrations for both induced and spontaneous mutants also did not differ between species for either haploids or diploids (chi-square test, p > 0.60 in all cases).

**Figure 1.**
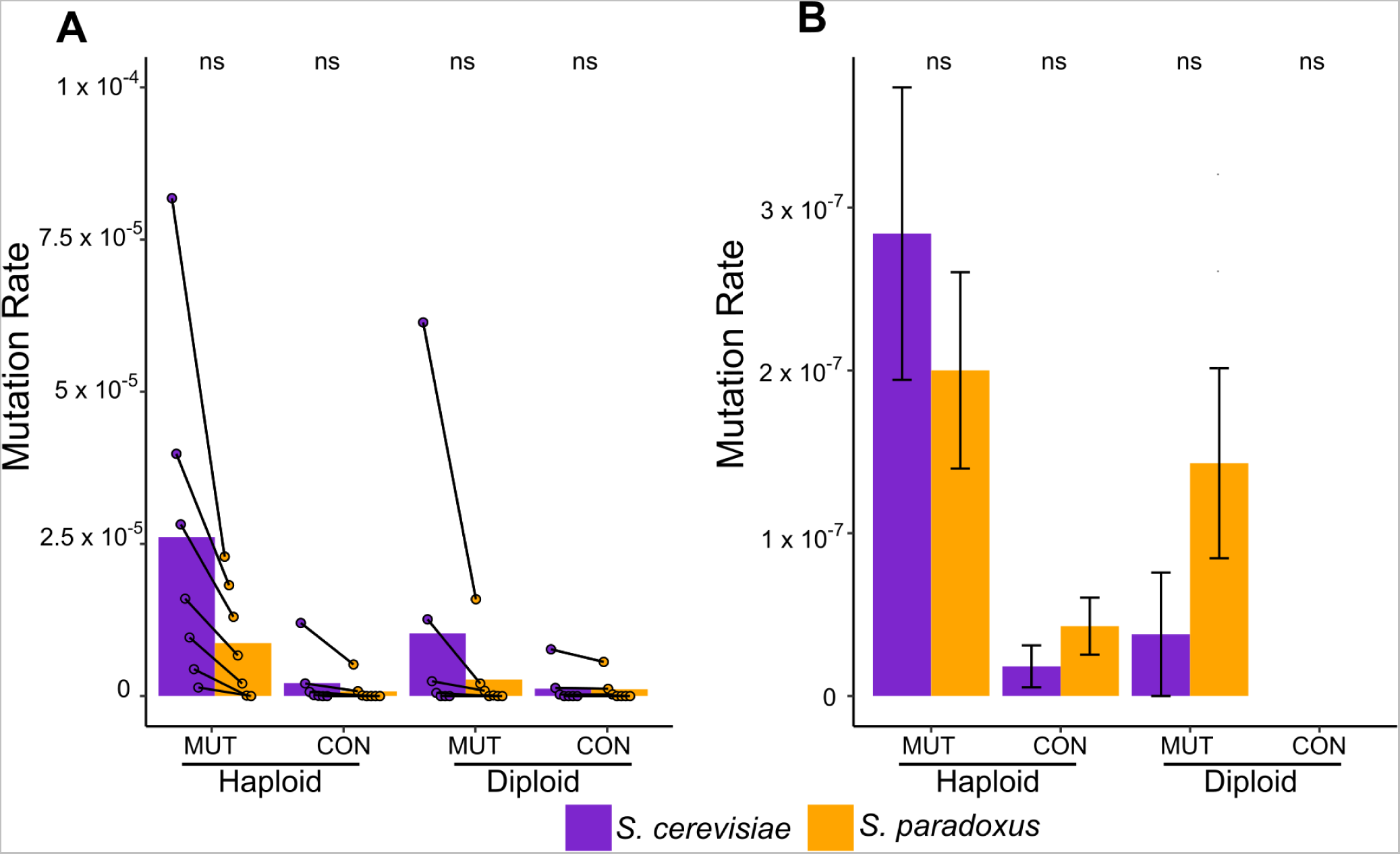
Mutation rate to copper or sulfite resistance does not differ between species. Mutation rates in platings of mutagenized and mock mutagenized pools of *S. cerevisiae* and *S. paradoxus*. (A) Copper mutation rates for haploids and diploids as measured by the number of colonies divided by the number of cells plated. Points represent individual measurements for different copper concentrations and are paired by concentration. Left to right, points signify mean mutation rate for the two ancestor strains on 0.1, 0.15, 0.2, 0.3, 0.4, 0.5, and 0.6 mM CuSO_4_. Bars signify overall mutation rate with all concentrations pooled together. (B) Sulfite mutation rates for haploids and diploids measured as the total number of non-escapee strains recovered over the total number of cells plated across all concentrations. Error bars represent the standard deviation of the mutation rate measurement assuming a Poisson distribution for colony counts. In both panels, mutagenized pool measurements are denoted as “MUT” and mock mutagenized pool measurements are denoted as “CON”. Significance between species was assessed via paired t-tests and chi-square tests where applicable, and “ns” denotes a nonsignificant difference.

At low sulfite concentrations we observed far more *S. cerevisiae* colonies than *S. paradoxus* colonies (Table S4). However, upon phenotyping we found that the vast majority of these “mutants” did not have a heritable sulfite resistance phenotype (see Methods). We defined these isolates as “physiological escapees”. After removing escapees, we retained only 31 haploid and 7 diploid mutants, and there were no differences in rates of induced or spontaneous mutants between species for either haploids or diploids (chi-square tests, p > 0.15 in all cases).

Taken together, the results from our initial mutant screen are not consistent with the target size hypothesis. We find that our controls indicate equal mutagenesis between species and that *S. paradoxus* haploids and diploids produce mutants with elevated copper and sulfite resistance at rates statistically indistinguishable from their *S. cerevisiae* counterparts. The species were also quite similar in the distribution of mutants recovered on the various copper concentrations tested.

### Mutant Effect Sizes

To test whether mutational effect size among mutations conferring elevated resistance to copper and sulfite differs between *S. cerevisiae* and *S. paradoxus*, we subjected a large subset of the mutants identified in our screen to a high throughput, robotics-based phenotyping assay. Briefly, we arrayed our mutants (N = 3,024 for haploids and N = 720 for diploids split evenly between copper and sulfite) onto solid plates containing many different concentrations of copper and sulfite. Our subset contained representative mutants from both the mutagenized and mock mutagenized pools isolated at various concentrations of copper and sulfite. We then quantified resistance by AUC (trapezoid rule) when colony sizes were plotted against stressor concentration (Figure S1B). Specifically, we measured the change in AUC relative to the ancestor (ΔAUC) and only retained for analysis those mutants whose AUC was three standard deviations higher than the ancestral measurements to exclude physiological escapees. Following escapee exclusion, the final sample sizes for haploid and diploid copper mutants were 1,107 and 309 respectively. For sulfite mutants these numbers were 31 and 7 respectively. These numbers represented total retention of 75.6% of the copper phenotype data and only 2.0% of the sulfite phenotype data due to the high number of physiological escapees.

We find that haploid copper mutants of *S. paradoxus* have a larger average effect size than *S. cerevisiae* copper mutants (Fig 2A). This result is driven by induced mutants (Kruskal-Wallis test, p = 4.7 x 10^-12^), and there is no species difference when only spontaneous mutants are considered (Kruskal-Wallis test, p = 0.06). In contrast, diploid copper mutants show no such species difference in effect size (Figure 2B, p = 0.25). There is also no species difference when these mutants are divided into induced and spontaneous and are considered separately (p = 0.08 and p = 0.12 respectively, Kruskal-Wallis test). Of note, YJF5538 the S288c derivative used as a control, displayed the maximum AUC possible in this assay of 0.8, indicating its copper resistance is far greater than that of any of the recovered mutants.

**Figure 2.**
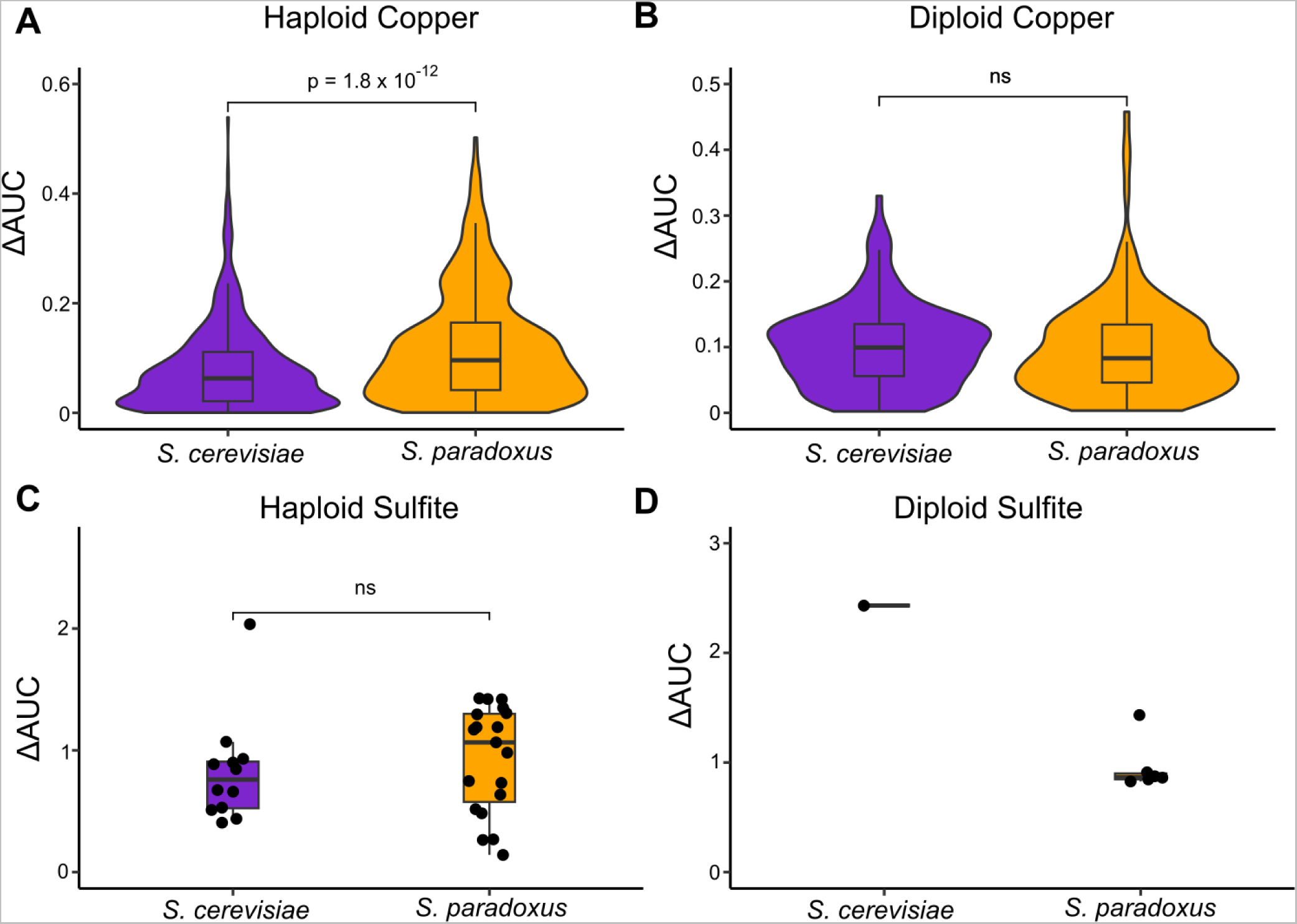
Limited species differences in effect size for copper and sulfite resistance mutants. Resistance is measured by ΔAUC of colony size as a function of stressor concentration in mM. Significance values are derived from Kruskal-Wallis tests (ns indicates not significant). (A) Haploid non-escapee copper mutants (N = 1,107) show a larger effect size in *S. paradoxus*. (B) Diploid non-escapee copper mutants (N = 309) show no species difference in effect size. (C) Haploid sulfite mutants (N = 31) show no difference in effect size. (D) Diploid sulfite mutants (N = 7) have a low sample size that precluded a high confidence comparison of effect size.

For sulfite we observed no difference in effect size between mutants of the two species among haploids (Figure 2C, p = 0.29, Kruskal-Wallis test). For diploids, we only recovered a single *S. cerevisiae* mutant that was not an escapee. Although this single ΔAUC measure was higher than the distribution of the six ΔAUC measures from the corresponding *S. paradoxus* mutants (p < 2.2 x 10^-16^, one sample t-test), these small sample sizes precluded high confidence inference of an effect size distribution for diploid sulfite mutants in these species (Figure 2D). Overall, the phenotyping data do not support the effect size hypothesis as being a major driver of the natural pattern of differential adaptation seen between *S. cerevisiae* and *S. paradoxus*. The distributions of effect sizes between species are largely similar, and the one well supported difference (haploid copper) does not align with the natural pattern of apparent constraints.

### Pleiotropic costs

To test whether resistance mutations in the two species tended to come with different pleiotropic costs, we also measured colony size on six permissive conditions for all mutants (Figure 3). We measured costs as growth relative to the ancestor. For copper mutants, we find pervasive costs in both species and sparse support for the costs hypothesis when the two species are compared. Specifically, we find that haploid copper mutants derived from both species have costs in all permissive conditions (Figure 3A, one-sample t-test, Bonferroni corrected, p < 10^-14^ in all cases). These costs show a difference between species in two of the six permissive conditions. Namely, in MM + glycerol and YPG, *S. cerevisiae* haploid copper mutants on average incur less severe costs than their *S. paradoxus* counterparts (Figure 3A, Kruskal-Wallis test, Bonferroni corrected, p < 0.05 in both cases). Similar patterns were seen in diploid copper mutants. *S. cerevisiae* diploid copper mutants incur costs in all six permissive conditions and *S. paradoxus* mutants incur costs in four of the conditions (Figure 3B, one sample t-test, Bonferroni corrected p < 0.05 in all cases). *S. paradoxus* mutants demonstrated higher costs than *S. cerevisiae* in three of the conditions (Kruskal-Wallis test, Bonferroni corrected, p < 0.05 in all cases).

**Figure 3.**
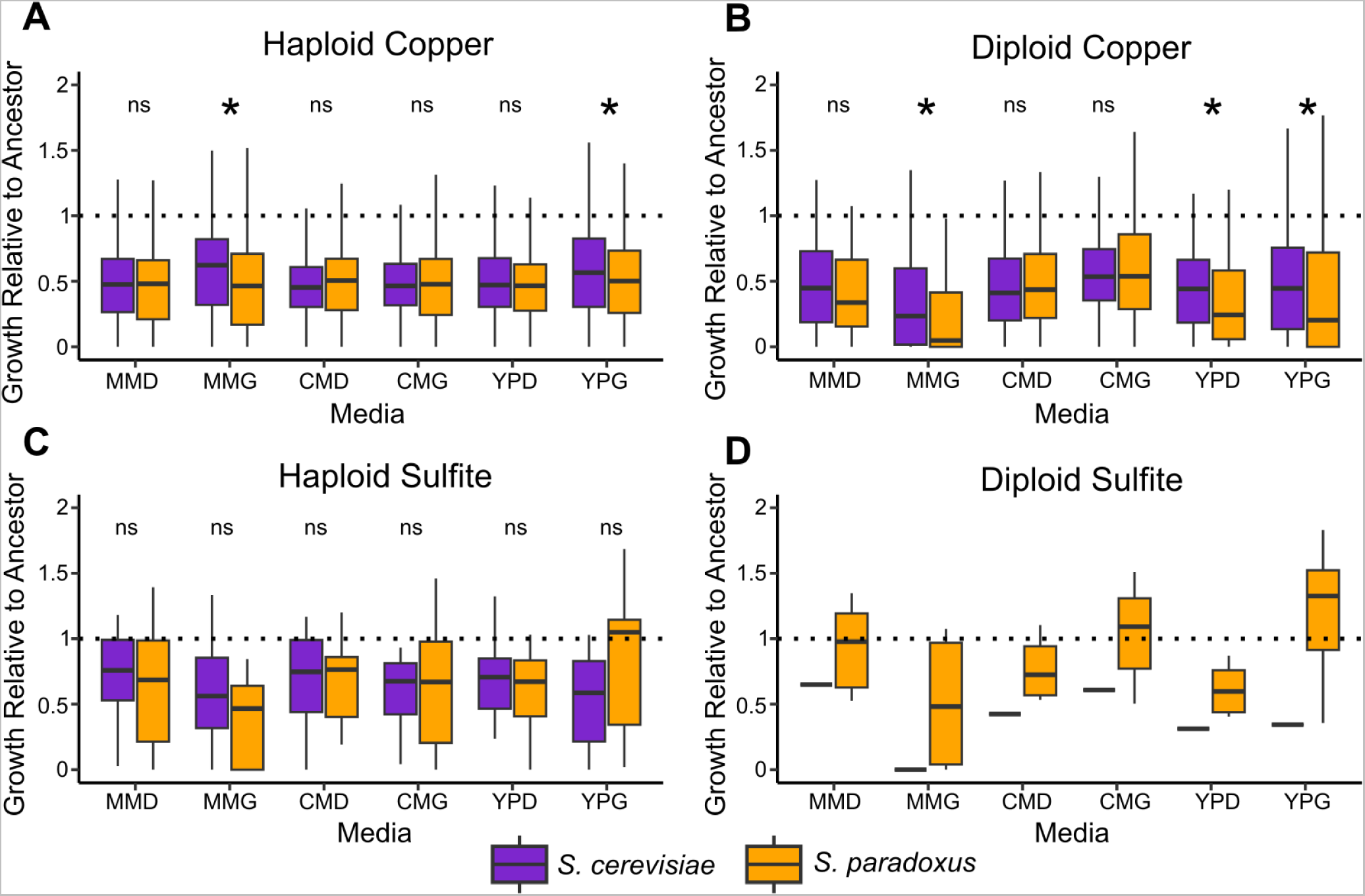
Species differences in pleiotropic costs of recovered mutations. Costs are measured by growth (colony size) in six media relative to the ancestor. Nonsignificant differences are denoted by “ns”, and significant differences are noted by an asterisk and represent p < 0.05 for a Kruskal-Wallis test after Bonferroni correcting for six comparisons. Media abbreviations are minimal media + dextrose (MMD), minimal media + glycerol (MMG), complete media + dextrose (CMD), complete media + glycerol (CMG), YP + dextrose (YPD), and YP + glycerol (YPG). (A) Costs for non-escapee haploid copper mutants with data in all permissive conditions (N = 999). *S. paradoxus* mutants display greater costs in MMG and YPG. (B) Costs for non-escapee diploid copper mutants with data in all permissive conditions (N = 286). *S. paradoxus* mutants display greater costs in MMG, YPD, and YPG. (C) Costs for non-escapee haploid sulfite mutants with data in all six conditions (N = 30). There are no significant differences in costs. (D) Costs for the diploid sulfite mutants recovered in this study (N = 7).

For haploid sulfite mutants, we observed costs among resistant mutants in four of six permissive conditions for *S. cerevisiae*, and in five of six permissive conditions for *S. paradoxus* (Figure 3C, one sample t-test, Bonferroni corrected, p < 0.05 in all cases). For *S. cerevisiae*, no costs were detected for sulfite mutants in MM + dextrose or CM + dextrose. For *S. paradoxus*, no costs were detected in YPG. Across the six conditions there were no differences in costs between species for sulfite mutants even before correcting for multiple comparisons (Kruskal-Wallis test, p > 0.05 in all cases). The single recovered *S. cerevisiae* sulfite mutant showed consistent costs across all six conditions. The six diploid *S. paradoxus* sulfite mutants only incurred costs in YPD relative to their ancestor (Figure 3D, one sample t-test, Bonferroni corrected, p < 0.05).

Following this, we also investigated the general relationship between costs and effect size across all mutants. Haploid and diploid copper mutants of both species showed a negative correlation between ΔAUC and cost relative to their ancestor (Table S11, Figure S5, Spearman rank correlation, p < 10^-5^ in all cases). Thus, mutants with larger effects on copper resistance tended to incur greater costs in both species. There was no interaction between species and resistance in predicting costs, indicating the relationship is similar for the two species (two-way ANOVA, p > 0.30 in both cases).

Haploid sulfite mutants showed a different pattern (Table S11, Figure S5) with no relationship between effect size and growth relative to the ancestor in *S. cerevisiae* (Spearman rank correlation, p = 0.19) but a negative relationship in *S. paradoxus* (Spearman rank correlation, p = 0.04). For diploid sulfite mutants, there was no relationship between effect size and costs for the six diploid *S. paradoxus* mutants. These results indicate that copper resistance mutations in both species tend to be more costly as their effects increase and that this relationship is less consistent across species for sulfite.

### Copper and sulfite resistance are caused by mutations in a small number of genes

To determine which mutations underlie copper and sulfite resistance in *S. cerevisiae* and *S. paradoxus*, we subjected 150 mutants and our four ancestral strains to whole genome sequencing (Table S3, N = 119 and N = 31 for copper and sulfite respectively). After filtering we found an average of 6.85 SNPs and insertions/deletions (indels) in each mutant. We found 20 genes with multiple disruptive variants within our dataset. Of these, nine were significant when compared to a simulated null distribution after correcting for multiple comparisons (see Methods). Of the nine, the three most commonly mutated genes were *PMA1* (copper, N = 42), *REG1* (copper, N = 11), and *RTS1* (sulfite, N = 9). Whereas *PMA1* and *RTS1* were mutated in both species, *REG1* mutations were only found in *S. cerevisiae*. There was no statistical difference in incidence of *PMA1* or *RTS1* mutants between species (chi-square test, p > 0.20 in both cases), but there was a difference in incidence between species for *REG1* mutations (chi-square test, p = 0.0013, Table S7). Of the remaining genes with multiple mutations, several were annotated as dubious ORFs or telomeric, indicative of genotyping errors rather than their having a role in copper or sulfite resistance.

*PMA1* mutations were called in 22 *S. cerevisiae* copper mutants and 20 *S. paradoxus* copper mutants. There was also one additional strain (YJF4439) not assigned a causal variant that harbored no coding changes in *PMA1* but had six synonymous changes in this gene. Among the 42 mutants with coding changes, there were six cases of site level parallel changes between species and there were no *PMA1* mutations detected among the sequenced sulfite mutants. This high level of incidence and parallelism shows that mutations in this gene cause elevated copper resistance in both species. It also lends support to the estimate of the saturation of the screen being greater than one mutation per site. *REG1* was mutated in 11 of the sequenced *S. cerevisiae* copper mutants, and no *REG1* mutations were found in *S. paradoxus* copper mutants. Of these, eight were nonsense mutations and two were frameshift mutations, signaling that loss of function *REG1* mutations confer copper resistance in *S cerevisiae*.

Among sulfite mutants, there was only one gene with a significant number of mutations: *RTS1*. This gene was mutated in five *S. cerevisiae* and four *S. paradoxus* sulfite mutants. Of these mutations, five were nonsense mutations, two were frameshifts, and two were missense mutations. These data indicate that loss of function mutations in *RTS1* in both species confer sulfite resistance. In addition to this gene, there was another gene that did not meet our significance criteria but showed a high degree of interspecies parallelism among the sulfite mutants, signaling that it is likely causal: *KSP1*. This gene is mutated in three sulfite resistant mutants of both species, representing a significant hit if the *S. cerevisiae* and *S. paradoxus* screens are considered together (p = 5.2 x 10^-5^, Bonferroni corrected). Of these six mutations, five of them are nonsense mutations, again signaling that loss of function mutations in this gene confer sulfite resistance in both species. Upon closer inspection of these mutants, it is likely that the three *S. paradoxus* mutants have a single origin, with each strain having exactly the same mutation and having been recovered from the same plate (Table S3).

For mutations in *PMA1*, *REG1*, *RTS1*, and *KSP1* we investigated their location and frequency across the gene body (Figures S6 and S7). We find that *PMA1* mutations in both species are dispersed broadly across the gene with more mutations being found at the C-terminus than the N-terminus. *REG1* mutations showed a strong N-terminal bias including a nonsense mutation at Lys12, indicating some of these mutations likely result in null alleles. For *RTS1*, recovered mutations were dispersed across the gene body, and for *KSP1* we saw N-terminal nonsense mutations in *S. cerevisiae*, with the one *S. paradoxus* mutation being found near the center of the gene body.

### Aneuploidy and CUP1 duplication cause copper resistance

Among the 119 copper mutants we sequenced, we found 45 strains that were aneuploid, nine of which were *S. cerevisiae* and 36 of which were *S. paradoxus*. Among copper mutants, almost every instance of aneuploidy either involved chromosome VIII, chromosome III, or both (Table S3). Based on a simulated null distribution, aneuploidy of chromosome VIII is enriched in both species (p < 5 x 10^-6^ in both cases) and aneuploidy of chromosome III is enriched in *S. paradoxus* (p < 10^-7^) but not *S. cerevisiae* (p = 0.36). This indicates that chromosome VIII aneuploidy is contributing to copper resistance in both species, and chromosome III aneuploidy is contributing to copper resistance in *S. paradoxus*. Although several *S. paradoxus* and *S. cerevisiae* strains have extra copies of both chromosome VIII and III, the frequency of this combination did not suggest an excess of co-occurrence given the frequencies of each respective aneuploidy (p > 0.5 in both cases). Comparing the two species, incidence of aneuploidy of any kind, along with aneuploidy of either chromosomes III or VIII were all significantly higher in *S. paradoxus* copper mutants (chi square test, p < 0.01 in all cases, Table S7). The *CUP1* gene resides on chromosome VIII and likely explains aneuploidy-based copper resistance.

We also called structural variants in our sequenced strains using Delly. We only found five cases of high confidence structural variants in our sequenced strains, and none could be assigned as likely causal for phenotype (Table S3, Table S5). Many prior studies have seen that expansions of *CUP1* tandem arrays confer greater copper resistance (Fogel et al. 1983, Adamo et al. 2012, Gerstein et al. 2015). To ensure *CUP1* duplications were not missed, we manually inspected coverage at the *CUP1* locus for each sequenced copper resistant strain. We found evidence of duplication of this gene in one *S. paradoxus* strain, YJF4464, with coverage data supporting 9 *CUP1* copies (Figure 4). Following mapping to a modified reference with an inverted repeat of the *CUP1* region, we find reads which span the repeat junction and support an inverted repeat structure. The duplicated region also is flanked on either side by two pairs of short, inverted repeats, suggesting the possibility of Origin-Dependent Inverted-Repeat Amplification (ODIRA) as the mechanism of this mutation (Brewer et al. 2011). We also manually inspected the *SSU1* locus in the sulfite mutants and found no anomalous patterns in any of the strains.

**Figure 4.**
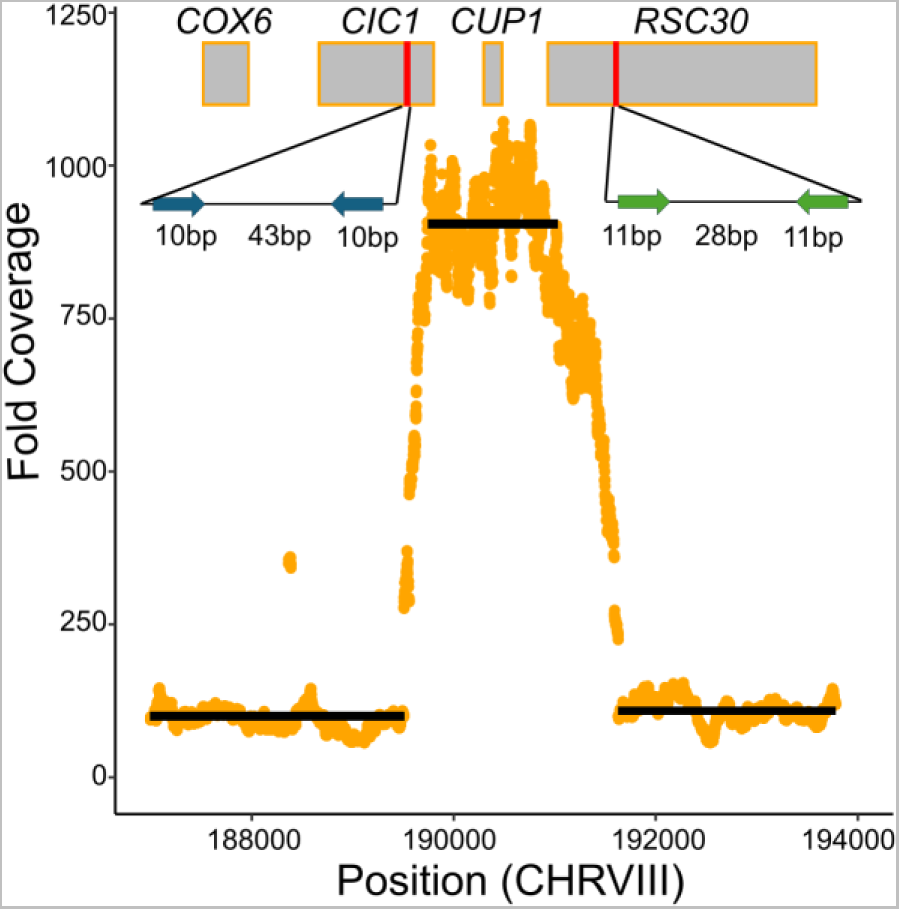
A *de novo CUP1* duplication occurred in an *S. paradoxus* mutant. The plot shows fold coverage as a function of chromosome VIII position for the *S. paradoxus* copper mutant YJF4464. Each point is a single nucleotide in the reference genome. Black lines represent average coverage across the positions they span. Rectangles are positions of genes plotted to scale. Coverage in this strain supports approximately 9 copies of *CUP1*. Red vertical segments represent two regions containing short, inverted repeats in the *S. paradoxus* genome. The insets are schematic depictions of these regions with the lengths of the repeats and intervening sequences. Inset colors indicate that the two regions contain repeats that are not identical in sequence to one another. Insets are not drawn to scale.

### Significant variants: effect size and costs

We performed several comparisons between species for all of the inferred causal variants, which were *PMA1*, *REG1*, *RTS1*, *KSP1*, chromosome VIII aneuploidy, chromosome III aneuploidy, and *CUP1* duplication. For all variants that had representative mutants derived from both species, we compared their effect size and costs in permissive conditions. We also performed these comparisons for the subset of strains that had not been assigned a causal variant.

When comparing effect size for similar mutations between species (Figure 5A and 5B), we find that there is only one significant difference. Mutations in *RTS1* tended to be of larger effect in *S. paradoxus* (Kruskal-Wallis test, p = 0.014). All other comparisons of effect size including mutants with unknown causal variants were not significant (Kruskal-Wallis test, p > 0.08 in all cases). We find that the *CUP1* duplication strain has a ΔAUC of 0.256, placing it among the more resistant mutants. When costs were compared within mutant classes (Figure 5C and 5D), we found that average costs relative to the ancestor in the six permissive conditions only significantly differed between species for *PMA1* mutants such that *S. cerevisiae PMA1* mutants incur less severe costs than their *S. paradoxus* counterparts (Kruskal-Wallis test, p = 0.01). All other comparisons were not significant (Kruskal-Wallis test, p > 0.08 in all cases). The *CUP1* duplication strain incurred an average cost relative to its ancestor of 0.54 across the six permissive conditions, indicating substantial costs.

**Figure 5.**
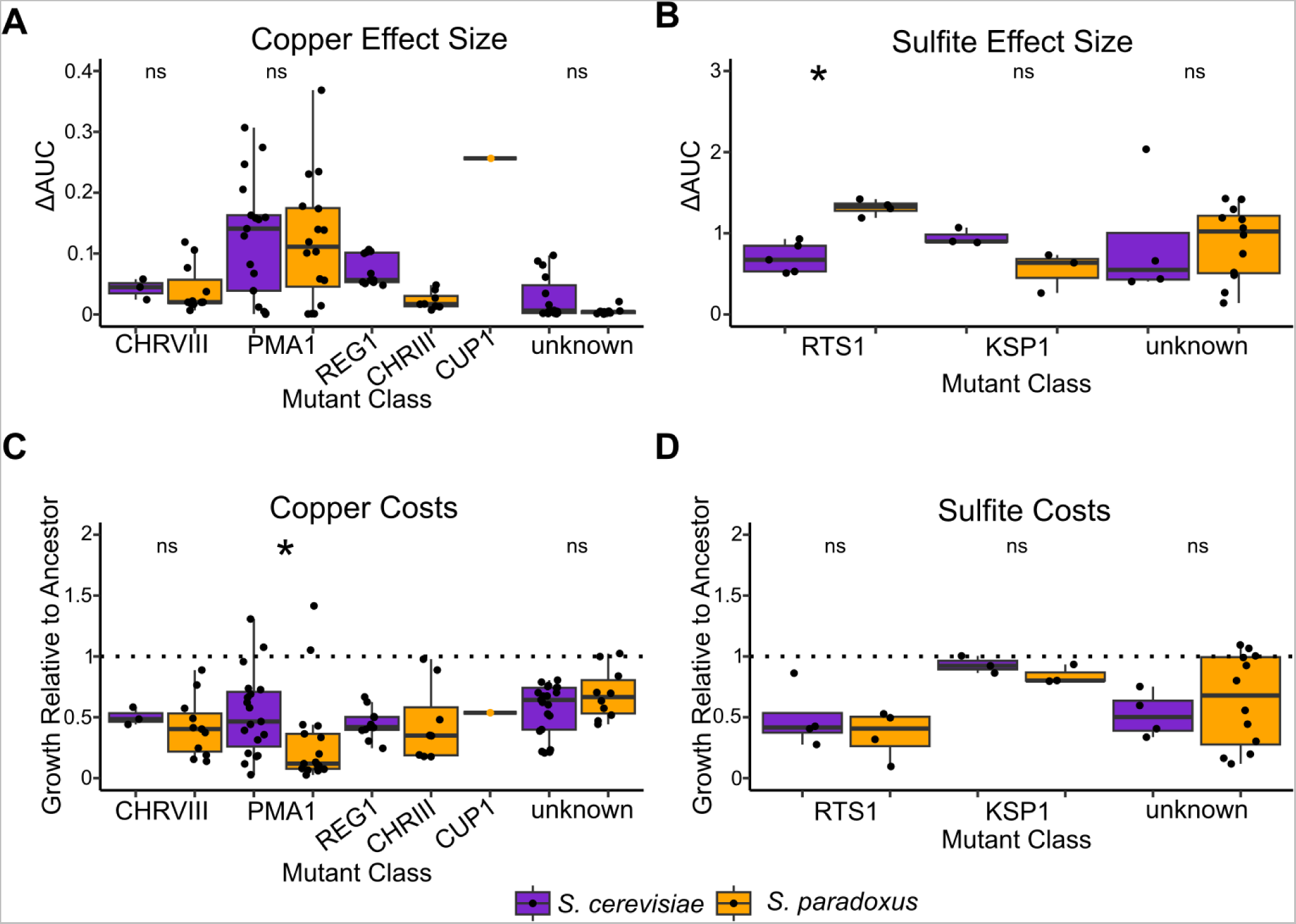
Effect sizes and costs for the mutant classes assigned as causal. Effect sizes are measured as ΔAUC for the colony size measurements as a function of stressor concentration in mM. Costs are measured as average growth relative to the ancestor across six permissive conditions. Significant species differences are noted with an asterisk and represent p < 0.05 for a Kruskal-Wallis test after a Bonferroni correction, and “ns” denotes a nonsignificant difference. (A) Copper effect sizes for the major copper mutant classes recovered for both species. There are no significant species differences. (B) Sulfite effect sizes for the major sulfite mutant classes recovered for both species. *RTS1* mutants have a significantly higher effect size in *S. paradoxus*. (C) Costs for the major copper mutant classes. *PMA1* mutants incur greater costs in *S. paradoxus*. (D) Costs for the major sulfite mutant classes. There are no significant species differences.

### The PMA1 copper resistance mutations are partially recessive loss of function mutations

*PMA1* encodes a membrane bound P2-type ATPase that exports protons from the cell to regulate intracellular pH and to establish an electrochemical gradient for secondary active transport into the cell (Young et al. 2023). Loss of function mutations in *PMA1* cause growth defects, pH sensitivity, sensitivity to cationic drugs, and have variable dominance (Morsomme et al. 2000, Cyert and Philpott 2013). If a weaker proton gradient mediates copper resistance in *PMA1* mutants, we expect our mutants to show pH sensitivity, a hallmark of loss of function mutations in *PMA1* (Cyert and Philpott 2013). To test if recovered *PMA1* mutants are loss of function mutants, we performed spot dilutions of 39 of our mutants onto MM and low pH MM (pH = 2.5). We found that 27 of the 39 mutants tested showed a clear low pH sensitivity phenotype (Table S8). When sensitivity is compared with copper resistance, we find that for both species low pH sensitive *PMA1* mutants are more copper resistant than mutants that are not sensitive to low pH media (Kruskal-Wallis test, p < 0.007 in both cases). This indicates that the greater the magnitude of loss of function in Pma1, the greater the copper resistance.

We also investigated the dominance of *PMA1* mutations (Table S9). To test whether our recovered mutations are dominant or recessive, we backcrossed five mutants with site level parallelism for both species to their ancestors yielding 10 heterozygous diploid strains. We find that in 7/10 cases, heterozygotes display intermediate sensitivity to low pH MM, and we find in 10/10 cases, the heterozygotes have intermediate copper resistance compared to their ancestors. In contrast, 8/10 diploids showed little or no cost on CM plates (Table S9). The results for the mutation Ala506Val are shown in Figure 6 for both species. This mutation displays incomplete dominance regarding low pH sensitivity and copper resistance in *S. cerevisiae* and *S. paradoxus*. These data point to a general pattern of incomplete dominance for *PMA1* mutations in both species.

**Figure 6.**
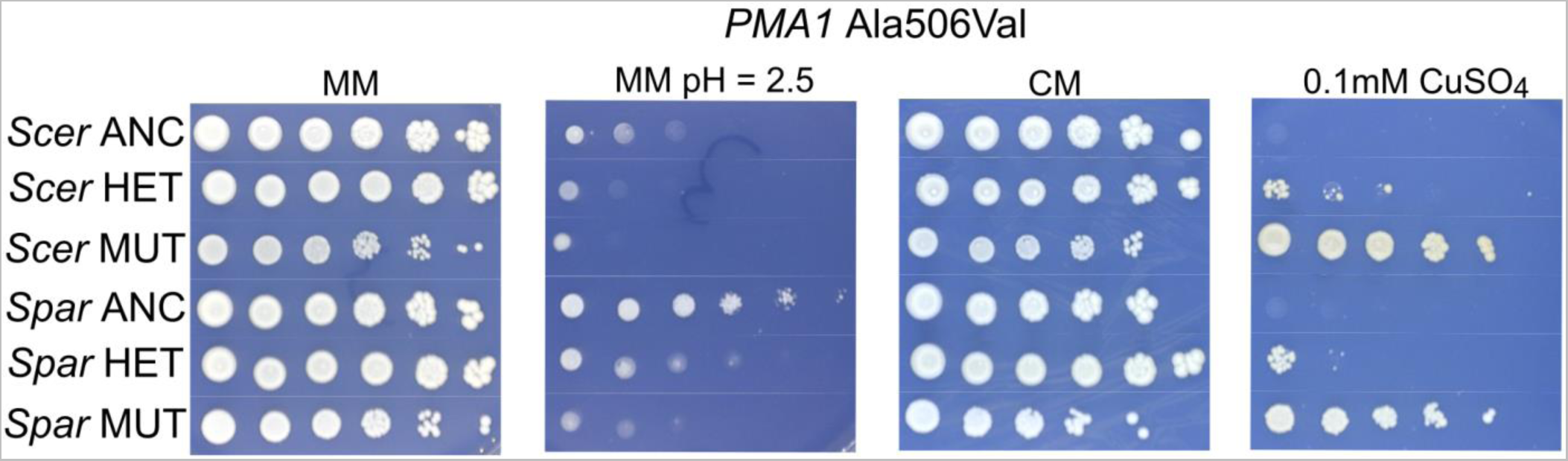
Spot dilution phenotypes associated with the Ala506Val mutation in *PMA1* in *S. cerevisiae* and *S. paradoxus*. *S. cerevisiae* is denoted as “*Scer*”, and *S. paradoxus* is denoted as “*Spar*”. Haploid ancestors (ANC) are tolerant of low pH media and sensitive to 0.1 mM CuSO_4_. Backcrossed heterozygotes (HET) show intermediate low pH tolerance and copper resistance compared to their ancestors and the haploid mutants. Haploid mutants (MUT) display low pH sensitivity and copper resistance.

### The REG1 deletion phenotype is species dependent

The recovery of *REG1* mutations in *S. cerevisiae* but not *S. paradoxus* led us to hypothesize that null alleles of *REG1* have a copper resistance phenotype in *S. cerevisiae* but not *S. paradoxus*. Additionally, some prior literature has drawn functional connections between *REG1* and *PMA1* through the action of Reg1 on the Glc7 complex, which can play a regulatory role in Pma1’s activity (Willaims-Hart et al. 2002, Cyert and Philpott 2013, Mazón et al. 2015, Guarini et al. 2024). This also led to the hypothesis that Reg1’s effect on copper resistance in *S. cerevisiae* may be mediated through an effect on Pma1 activity.

In the context of these hypotheses, we performed two follow up experiments related to *REG1*. First, we spot diluted several *REG1* nonsense mutants onto low pH MM to determine if these mutants, like the *PMA1* mutants, were low pH sensitive. Second, we deleted *REG1* from one of the *S. cerevisiae* ancestral strains (YJF3732) and one the *S. paradoxus* ancestral strains (YJF3734) to determine if the effect of inactivating *REG1* varied between species. In the first experiment, we found that the *REG1* nonsense mutants we assayed have a low pH sensitive phenotype, phenocopying *PMA1* mutants (Figure 7A). This is consistent with the hypothesis that the copper resistance effect of these mutations may be mediated through an effect on Pma1. In the second experiment, we tested the copper resistance and low pH tolerance of deletion mutants of *S. cerevisiae* and *S. paradoxus*. The *S. paradoxus* deletion mutant had an extreme slow growth phenotype upon isolation. As such, we also isolated a suppressor mutant derived from this deletion strain to include in our phenotype assay, and we included a 10x concentrate of the original mutant in the assay as well (Figure 7B and 7C). We find that deletion of *REG1* in *S. cerevisiae* has a small effect on growth for CM and MM. However, in *S. paradoxus* there is a strong growth defect in these permissive conditions. Wild type growth rates in nonstress conditions were restored in the *S. paradoxus* suppressor mutant. For both species, deletion of *REG1* leads to increases in copper tolerance. However, this gain appears to be slightly greater in *S. cerevisiae*. A low pH sensitive phenotype is apparent in both deletion strains but not the *S. paradoxus* suppressor mutant. These data are consistent with diverged mutational effects at the *REG1* locus between species and suggest *REG1* mutations may not be a viable route to copper adaptation in *S. paradoxus* despite an effect on copper tolerance being conserved. They also provide indication that Reg1’s effect on copper tolerance may be mediated through an effect on Pma1 activity.

**Figure 7.**
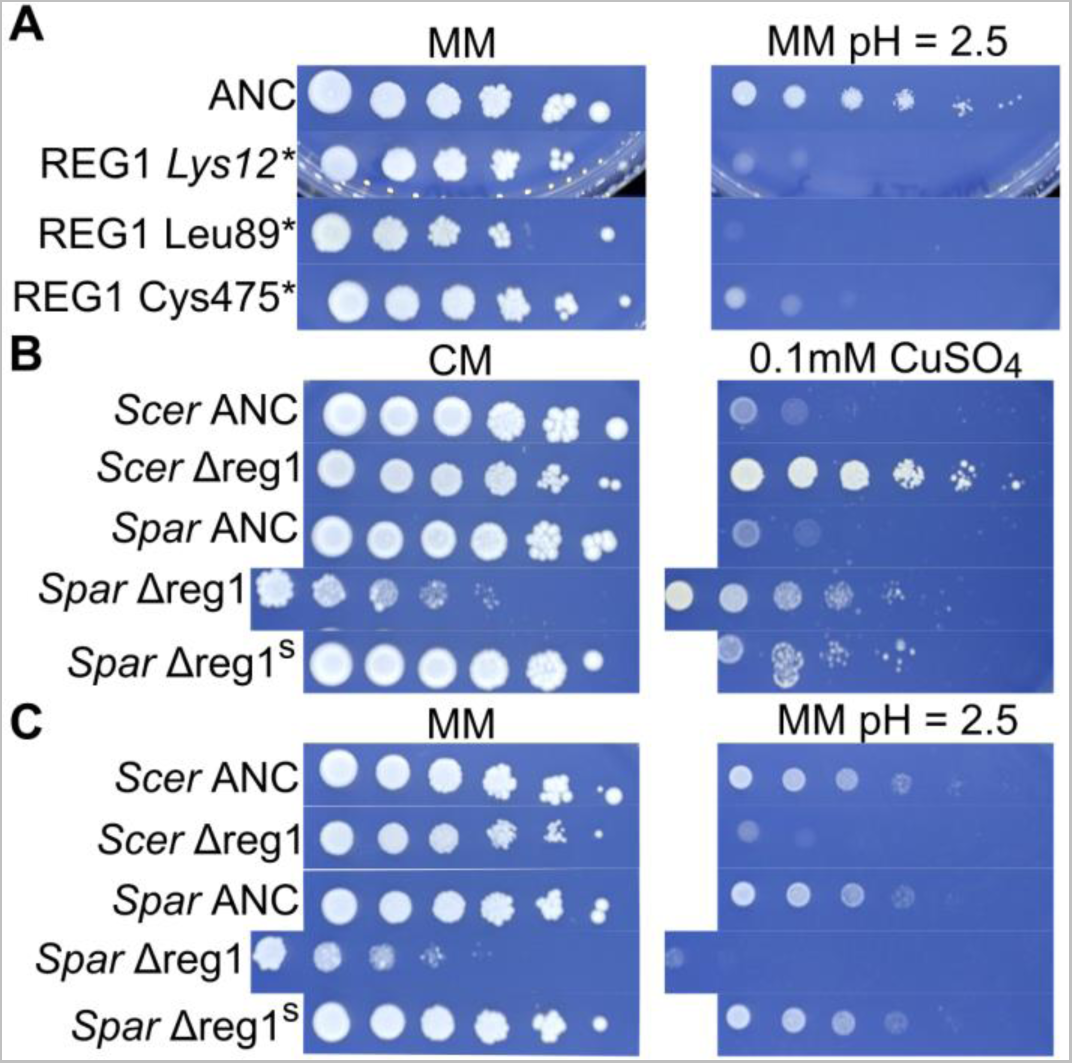
Spot dilution phenotypes associated with *REG1* loss of function in both species. (A) *S. cerevisiae* strain YJF3732 (ANC) and three *REG1* nonsense mutants identified among sequenced strains. All three mutants show low pH sensitivity. (B) *S. cerevisiae* strain ancestor (YJF3732) and *S. paradoxus* strain ancestor (YJF3734) along with *REG1* deletion strains derived from these strains. *Spar* Δreg1^s^ refers to a suppressor mutant derived from “*Spar* Δreg1”. Deletion of *REG1* leads to a severe growth defect in *S. paradoxus* and confers elevated copper resistance in both species. (C) Deletion of *REG1* leads to low pH sensitivity in *S. cerevisiae* and *S. paradoxus*. However, the suppressor mutation rescues wild type growth.

## Discussion

In nature, *S. cerevisiae* and *S. paradoxus* show contrasting patterns of adaptation regarding the anthropogenic enological stressors copper and sulfite. Although both species are present in vineyards, only *S. cerevisiae* has adapted to these chemicals, and it has done so multiple times. In this study, we investigated divergence in the DME for copper and sulfite resistance. Overall, our results show broadly conserved mutational effects along with high levels of gene-level and even site-level parallelism between species. However, we also found the subsets of mutations conferring resistance were not identical for the two species and that mutations in homologous genes could display species specific effect sizes and costs. Broadly, these results do not provide a clear explanation for the absence of copper and sulfite resistance in *S. paradoxus*, but they highlight the presence of differences in the DME between species and some of the limitations of experimental characterization of the DME.

### Similarities and differences in the DME

In our first set of experiments, we found that the mutational target size for copper and sulfite resistance was very similar between species, reflecting minimal divergence in terms of mutational availability concerning adaptation to these stressors. However, our assessment of the mutational target size differences for sulfite resistance was limited by the large number of physiological escapees. There are several potential explanations for this including stochastic variation in expression of *SSU1*. That said, the most plausible explanation is likely technical rather than biological. Sulfite plates have long been known to be difficult to work with due to off gassing of SO_2_ (Park et al. 1999), which can lead to variable sulfite concentrations batch to batch, and perhaps even within individual plates. We addressed the technical challenge of pouring sulfite containing agar by adding sodium sulfite to 50 mL aliquots of media and saw relative success in obtaining uniform toxicity in our plates. However, this method was likely still imperfect, and given the large number of cells we screened, some were able to produce colonies during the initial plating despite having no mutations conferring greater resistance.

After we screened for mutants, we phenotyped a large number of them and assayed their effect size. Similar to target size, distributions of effect sizes for *S. cerevisiae* and *S. paradoxus* were largely similar. There was one significant difference between species which was that haploid *S. paradoxus* copper mutants had a larger average effect size than their *S. cerevisiae* counterparts. In the context of the sequencing results, there is not an obvious explanation for the difference between haploid copper mutants of the two species. Within the main classes of copper mutants we observed (chromosome VIII aneuploids and *PMA1* mutations), we did not see a difference in effect size between species. However, this subset is biased for the most common types of mutations that confer copper resistance. Thus, it appears likely that rarer mutant classes that we were unable to detect in our sample underlie the species difference in effect size we see for haploid copper mutants.

Additionally, the species difference for copper does not persist in diploids, signaling the possibility of differences in dominance of copper resistance mutations between species. The most common type of mutation recovered for copper was loss of function mutations in *PMA1*. When backcrossed to the ancestral strain, these mutations showed a consistent pattern of incomplete dominance in both species, signaling that differences in dominance for *PMA1* mutations likely do not explain the difference between the haploid and diploid copper results. Variable effects of aneuploidy on haploids and diploids are expected due to differences in the relative increase in gene dosage. If these variable effects also differ between species, it remains a possibility that aneuploidy could account for some of the effect size differences between haploids and diploids.

Regarding sulfite, effect sizes were not different between species. However, in the context of the sequencing results, mutations in *RTS1* were present in both species, but their effect sizes differed significantly. This illustrates an interesting property that can be displayed by the DME: Although the DME can overall be quite similar between species, there can still be meaningful gene-level idiosyncrasies in effect size despite these general similarities.

Following the effect size assay, we investigated the pleiotropic costs of our mutants. We measured costs using growth in permissive conditions relative to the ancestor. We found that *S. paradoxus* mutants with elevated copper resistance tended to incur greater costs than their *S. cerevisiae* counterparts in several permissive conditions for both haploids and diploids. We found that this difference is not explained by differences in costs between strains aneuploid for chromosome VIII. In contrast, we do observe significantly greater costs in *S. paradoxus PMA1* mutants compared to their *S. cerevisiae* counterparts. This common mutant class may therefore partially explain the few differences in costs we do observe. Similar to effect size, the pattern may also in part be driven by rarer mutant classes that we were not able to identify in the sequencing data due to an insufficient sample size.

Taken at face value, the presence of greater costs in *S. paradoxus* in a few conditions offers an attractive potential explanation for apparent differences in adaptive capacity for copper. In principle, if such a difference were meaningful in nature it could increase the waiting time for a second beneficial mutation to occur before local extirpation, a phenomenon theoretically understood to increase the probability of evolutionary rescue in declining populations (Orr and Unkless 2014). However, there are many complications that can affect how pleiotropic costs impact adaptation in nature. For example, costs were pervasive in our data for both species, in contrast to the lack of apparent costs in natural copper and sulfite resistant *S. cerevisiae* strains. It could be that rare, costless mutations are the most relevant to adaptation. If this is the case, then a species difference in costs (among costly mutations) may not be ecologically relevant because these mutations never persist in nature for either species. Without a class of resistance mutations that are costly in *S. paradoxus* but not in *S. cerevisiae*, costs do not offer a clear explanation for natural outcomes.

Mutations that ameliorate costs are another complication that make it difficult to get a clear picture of how costs may affect adaptation in these species. It could be that costs are in fact pervasive for all mutations conferring copper and sulfite resistance including *SSU1* rearrangements and *CUP1* duplications, and the lack of costs seen in strains harboring these mutations is due to secondary mutations that compensate for the costs. If compensatory mutations play a role in adaptation to these stressors, the role of costs in determining evolutionary outcomes would be difficult to assay in a study of a single mutational step. Another possibility is that there are species differences in costs for very rare, large-effect mutations like *SSU1* rearrangements and *CUP1* duplications. We only observed a single *S. paradoxus* strain that harbored a *CUP1* duplication. This strain did exhibit costs in permissive conditions, in contrast to what is known about domesticated *S. cerevisiae* strains. However, with a sample size of one and without having a *de novo CUP1* duplication in *S. cerevisiae* as a point of comparison, there remain alternative possibilities. Duplication of *CUP1* could be costly in both species, but only *S. cerevisiae* can compensate for these costs while retaining resistance, or both species could have the capacity to compensate for the costs with secondary mutations. These many unknowns concerning how costs may affect adaptive trajectories limit the inferences that can be made from the species differences in costs we observe. Regardless of their relevance in nature, our results indicate that pleiotropic costs to resistance can be variable in different genetic backgrounds and species. Overall, the costs data trend in the direction expected under a model of differential mutational constraints, but they do not offer clear explanatory or predictive insight into the natural outcomes seen in vineyard strains of both species.

### Identity and effects of sequenced mutants

Among our sequenced strains, we were able to identify many causal variants underlying copper and sulfite resistance in both species. For copper, our screen yielded a large number of strains harboring mutations in the gene encoding the essential membrane bound proton efflux pump *PMA1*. The primary functions of Pma1 are to regulate intracellular pH, and to establish an electrochemical gradient for secondary active transport. Pma1 is biochemically well characterized (Morsomme et al. 2000, Young et al. 2024), and mutations in this gene have been implicated in copper resistance previously (Gerstein et al. 2015). The most plausible mechanism for this effect based on our experiments appears to be reduced copper influx due to a weaker electrochemical gradient (Morsomme et al. 2000, Cyert and Philpott 2013). This is consistent with other studies that have found loss of function mutations in *PMA1* that increase resistance to other cations and cationic drugs (Williams-Hart et al. 2002, Cyert and Philpott 2013). Thus, our results indicate that loss of function mutations in *PMA1* are viable mutational routes to greater copper resistance in *S. cerevisiae* and *S*. *paradoxus*. However, the ecological relevance of these mutations is highly questionable. Wine fermentations are typically very acidic environments and proper Pma1 function can be rate limiting for growth (Cyert and Philpott 2013, Williams et al. 2015). On these grounds we speculate that the costs of *PMA1* mutations are likely far too detrimental to allow for these mutations to persist in vineyards or contribute to domestication associated copper resistance.

The other major class of copper mutations shared by the two species was aneuploidy of chromosome VIII. For *S. paradoxus*, aneuploidy of chromosome III was also significant, but the reason for this phenotype is not obvious because none of the major genes implicated in copper resistance in yeast such as *CUP1*, *CUP2*, *CRS5*, and several others reside on this chromosome (Cullotta et al. 1994, van Backel et al. 2005, Gerstein et al. 2012, Caudy et al. 2013, Chang et al. 2013). The gene *HSP30* resides on chromosome III in both species, and this gene is a negative regulator of *PMA1*. It is plausible this gene may contribute to the copper resistance phenotype we see, but further work would be needed to confirm this. In contrast, the explanation for the phenotype of chromosome VIII aneuploidy is easily accounted for because the *CUP1* gene resides on chromosome VIII in both species. This aneuploidy has been known to cause copper resistance for many decades (Fogel et al. 1983) and was also seen in the study by Gerstein et al. (2015). We saw a significantly higher incidence of chromosome VIII aneuploidy in *S. paradoxus* compared to *S. cerevisiae*. However, effect size and costs were indistinguishable between species. Aneuploidy is caused by nondisjunction and occurs at appreciable rates in *S. cerevisiae* with higher rates for smaller chromosomes (Gilchrist and Stelkens 2019). It has been argued that aneuploidy represents an excellent avenue for short term adaptation to drastic changes to the environment because it often represents an accessible mutational path that can have large effects due to its increase in gene dosage for the aneuploid chromosome (Gerstein and Berman 2015). Aneuploidy has also been hypothesized to be valuable to adaptation because it is reversible. Indeed, Yona et al. (2012) observed that aneuploidy of chromosome III was adaptive under a high temperature selection scheme, but that after other beneficial mutations arose, the evolving strain eventually lost its extra chromosome. In this way, aneuploidy can be seen as a short-term adaptive solution that increases the potential waiting time for a rarer adaptive variant to arise.

Chromosome VIII aneuploidy is of particular interest for copper resistance due to how it may impact the rate *CUP1* tandem expansion. As pointed out by Zhao et al. (2014) and Fogel et al. (1983), tandem expansion of a multi-copy *CUP1* array is a fundamentally different process and far more common than *de novo* formation of a tandem array from a single copy. Many studies have observed expansion of *CUP1* arrays that were already multi-copy, and this occurs via unequal crossovers (Fogel and Welch 1982, Fogel et al. 1983, Adamo et al. 2012, Gerstein et al. 2015, Hull et al. 2017, Zhao et al. 2017). However, to our knowledge no prior study has explicitly observed *de novo* formation of a *CUP1* array from a strain that began with a single copy until now. Fogel et al. (1983) posited that aneuploidy of chromosome VIII may potentiate *CUP1* tandem expansion. This would align with the proposed mechanism of how *CUP1* tandem arrays originate, which requires two simultaneous double strand breaks on separate chromosomes and erroneous non-homologous end joining of the two chromosomes (Zhao et al. 2014). However, we did not observe any tandem duplications, and the repeat structure we observed is better explained by a different model.

Upon close inspection of the *CUP1* duplication in *S. paradoxus*, we found evidence for an inverted repeat structure similar to a *SUL1* inverted repeat structure described by Araya et al. (2010). As elaborated by Brewer et al. (2011), the Origin Dependent Inverted-Repeat Amplification (ODIRA) model offers a potential explanation for how these structures can arise. The model requires a nearby origin of replication and nearby flanking short, inverted repeats. Consistent with this model, the *CUP1* locus is nearby the *ARS810* locus in *S. cerevisiae*, and in *S. paradoxus* there exists a pair of short (10-11bp), inverted repeats directly upstream and downstream of the duplicated region we observe. Only *S. paradoxus* harbors inverted repeats of 10bp or more on either side of the *CUP1* region, but smaller inverted repeats (7bp or fewer) are highly abundant in both species on either side of the gene. The coverage pattern also increases and decreases in a stepwise pattern, similar to the CNVs described by Todd and Selmecki (2020).

If this duplication arose via this mechanism, it would raise many questions about the origin of *CUP1* duplications. Namely, this mechanism does not appear to be how the known *CUP1* duplications in domesticated strains of *S. cerevisiae* arose, so why have these structures not arisen in *S. cerevisiae*? One potential explanation is that *S. cerevisiae* lacks inverted repeats of sufficient length at this locus to drive the ODIRA mechanism. Irrespective of the explanation, our data demonstrate evidence for a different mechanism of *CUP1* amplification in *S. paradoxus* than what has been documented in *S. cerevisiae*.

The final set of copper resistance mutations we identified were loss of function mutations in *REG1* in *S. cerevisiae*. We did not observe any mutations in *REG1* in *S. paradoxus*, and this prompted us to delete this gene in both species. We found that deletion of *REG1* in *S. paradoxus* led to an extreme slow growth phenotype and increased copper resistance, indicating this growth defect precluded its detection in the mutant screen. Our results suggest that there has been functional divergence between *S. cerevisiae* and *S. paradoxus* in relation to Reg1 function, but the copper resistance phenotype caused by its absence is conserved between species. We also performed low pH sensitivity assays on *REG1* deletion mutants and mutants recovered in the screen. This was motivated by the fact that Reg1 is a regulatory subunit of the Glc7 complex which can play a role in regulating Pma1 activity (Willaims-Hart et al. 2002, Cyert and Philpott 2013, Mazón et al. 2015, Guarini et al. 2024). We find that all of the *REG1* mutants tested display a low pH sensitive phenotype consistent with the absence of functional Reg1 leading to increased copper resistance via lowering Pma1 activity. Importantly, the result does not exclude the possibility of Reg1 exerting an effect on low pH tolerance independent of Pma1, but this seems unlikely given that Pma1 is the primary regulator of cytosolic pH (Cyert and Philpott 2013). Curiously, the idea that lack of *REG1* may lower Pma1 activity appears to be overtly contradicted in a biochemical assay of Pma1 activity in a *REG1* deletion background (Young et al. 2010). Two other studies have also investigated Glc7’s capacity to dephosphorylate Ser899, Ser911, and Thr912 in a *REG1* deletion background and found these functions are not dependent on the presence of *REG1* (Mazón et al. 2015, Guarini et al. 2024). Thus, Reg1 and Pma1 appear to interact via the Glc7 complex, and our data point to the absence of functional Reg1 lowering Pma1 activity, but the precise functional relationship between these genes remains unclear. It is also unclear whether the higher cost of *PMA1* mutants in *S. paradoxus* is related to the higher cost of *REG1* deletions.

For sulfite, the main route to adaptation in domesticated *S. cerevisiae* strains has been via structural variants that alter the upstream sequence of *SSU1*. This type of mutation has occurred at least three times independently and includes two examples of reciprocal translocations and an inversion, all conferring similar levels of sulfite resistance (Goto-Yamamoto et al. 1998, Pérez-Ortın et al. 2002, Zimmer et al. 2014, García-Ríos et al. 2019). In many prior studies *FZF1* and *SSU1* have been the principal genes presumed to make the largest contributions to sulfite tolerance (Park et al. 1999). Surprisingly, we did not observe any mutations in these genes in our dataset. Instead, we found two causal genes, *RTS1* and *KSP1*, that have effects on sulfite tolerance in both species. *RTS1* encodes a regulatory subunit of protein phosphatase 2A, and it is thought to play several roles in mitosis (Shu et al. 1997). Null mutants of this gene have been shown to be heat sensitive, ethanol sensitive, and sensitive to several other drugs (Dudley et al. 2005, Yoshikawa et al. 2009). One interesting finding from Linderholm et al. (2008) points to a potential role in sulfite metabolism for *RTS1*. In this study, the deletion collection was screened for genes that affect the capacity to produce hydrogen sulfide. Hydrogen sulfide production is principally affected by the activity of sulfite reductase. In this screen, the *RTS1* deletion mutant was found to be a hyperproducer of hydrogen sulfide. In our data, we find that nonsense mutations in *RTS1* confer greater sulfite resistance. Taken together, these results suggest that null alleles of *RTS1* may exert an effect on sulfite tolerance via increasing the activity of sulfite reductase within cells. If true, this would represent a mechanistic departure from how sulfite tolerance has been achieved in domesticated strains, which universally increase sulfite efflux.

The other gene in which we observed disruptive hits in sulfite mutants for both species was *KSP1*. This gene encodes a serine/threonine protein kinase involved in TOR signaling and autophagy (Umekawa and Klionsky 2012). *KSP1* is also required for filamentous growth in *S. cerevisiae* (Bharucha et al 2008). Null alleles of this gene have been shown to increase resistance to a wide variety of chemicals including cycloheximide, fluconazole, and caffeine (Kapitzky et al. 2010). *KSP1* has not been explicitly implicated in sulfite tolerance to our knowledge. However, we speculate that this effect is likely mediated through its effect on autophagy. It was recently shown that autophagy is required for sulfite tolerance in *S. cerevisiae* (Valero et al. 2020), and Ksp1 is a negative regulator of autophagy (Umekawa and Klionsky 2012).

### Implications for copper and sulfite adaptation in nature

Natural domesticated isolates of *S. cerevisiae* that have adapted to resist copper and sulfite have done so via two known genetic mechanisms. For copper the mechanism is through tandem amplification of *CUP1* and for sulfite the mechanism is through rearrangements that alter the upstream sequence of *SSU1*, thereby increasing its expression. These mutations are of extremely large effect and would be easy to detect. None of our recovered mutants had copper resistance approaching that of YJF5538, an S288c derivative harboring multiple copies of *CUP1*. Additionally, based on the phenotypes obtained in prior studies, none of our sulfite mutants are as resistant as domesticated strains harboring rearrangements (Pérez-Ortın et al. 2002, García-Ríos et al. 2019). These comparisons support that large effect structural variations were not erroneously missed in our sample of sequenced strains. The apparent rarity of these mutations coupled with the fact that they have occurred in parallel in *S. cerevisiae* point to a strong likelihood that (short term) effective population sizes for this species in vineyards are likely extremely large (Karasov et al. 2010). However, *S. paradoxus* is likely comparably abundant in vineyards (Dashko et al. 2016). This raises a critical issue concerning studies of the DME in relation to adaptive outcomes in nature. Namely, extremely rare mutations that are beyond the detection limit of normal screens can play important roles in adaptation, especially for organisms with large population sizes like microbes. Measures of the DME may therefore be meaningfully biased for common mutations even if the screen is “saturated” for point mutations as ours was.

Although seemingly quite rare, we did recover a single strain with a mutation similar to those observed in domestication strains. The strain in question is an *S. paradoxus* strain that harbors approximately 9 copies of *CUP1*. This strain is fairly copper resistant and incurred consistent costs across the permissive conditions tested. These results are of interest because this strain has a modest copper resistance phenotype compared to YJF5538. The addition of 8 extra copies of *CUP1* would likely be expected to have a much larger effect in an *S. cerevisiae* background than we observed in *S. paradoxus* based on prior studies (Strope et al. 2015). This raises the possibility of differential effect size of *CUP1* mutations in the different species, but further investigation is needed especially considering the inverted repeat structure observed. Additionally, this strain’s persistent costs are inconsistent with what is known about copper resistant strains of *S. cerevisiae* harboring *CUP1* duplications, which have no apparent growth defect. As such, this result similarly raises the possibility of species-specific costs to *CUP1* amplification in *S. paradoxus*, but more detailed study is also needed.

These results raise many other questions about *CUP1* tandem arrays more generally. Firstly, it is unclear why duplication of this gene appears to be much rarer than for other genes such as *SUL1* (Sanchez et al. 2017). Close inspections of the known *CUP1* tandem structures have shown that the two genes flanking *CUP1*, *CIC1* and *RSC30*, are never included intact in the tandem duplications, which Zhao et al. (2014) noted as potentially indicating that extra doses of these genes are poorly tolerated. *CIC1* is also essential, meaning any *CUP1* duplication must keep a functional copy of *CIC1* intact. The *CUP1* duplicated region in *S. paradoxus* also only covers part of the *CIC1* and *RSC30* genes, similar to *CUP1* duplications in *S. cerevisiae*. These potential constraints based on genomic location may explain why this duplication seems to be far rarer than duplication of *SUL1* and other genes. Secondly, even if rare, *S. cerevisiae* and *S. paradoxus* are recovered at similar frequencies in vineyards and *S. cerevisiae* only becomes much higher abundance during fermentation (Dashko et al. 2016). Whether these species differ in the rate of this mutation or if the ecology of these species in vineyards differs more than is currently appreciated both remain open questions. What is clear is that *CUP1* duplications are relatively rare and may occur at a rate near the detection limit of the screen performed in this study. We screened ∼10^8^ mutagenized cells in total and recovered a single case of *de novo CUP1* duplication. Thus, the mutation rate for *CUP1* amplification may be on the order of 10^-8^, but with a sample size of only one the rate could be even lower.

Among the sulfite mutants we sequenced, we did not observe any rearrangements involving *SSU1*, meaning the rate of occurrence of these mutations is lower than the detection limit of the screen we performed. Translocations and inversions are relatively rare compared to other types of mutations, and it is conspicuous that these rare classes of mutants have repeatedly been selected for in sulfite exposed strains. This suggests that very few mutations of large effect are available to *S. cerevisiae* with respect to acquisition of greater sulfite resistance. Complicating the matter further, CRISPR induction of one of the translocations showed that this mutation lowers sulfite tolerance in a background lacking duplications of a 76 bp repeat in the *ECM34* promoter, demonstrating background dependent effects of this mutation (Pérez-Ortın et al. 2002, Fleiss et al. 2019).

Overall, these observations are consistent with the extremely small mutational target size we observed for sulfite in our mutant screen. It is likely that very few mutations in the genome aside from extremely rare rearrangements can elevate *SSU1* expression to the levels needed for success in a winemaking environment. This raises questions about the regulation of *SSU1* and the mutations available in its promoter sequence and its primary transcription factor, Fzf1. Namely, why is greater *SSU1* expression seemingly so mutationally inaccessible? *FZF1* has diverged considerably between *S. cerevisiae* and *S. paradoxus*, with the *S. paradoxus* allele actually being a more efficient activator of *SSU1* transcription (Engle and Fay 2012). In the context of this result, *FZF1* mutations that confer greater *SSU1* expression and sulfite tolerance may be expected to be more accessible in *S. cerevisiae* than *S. paradoxus*. The ecological relevance of *FZF1* and *SSU1* to these species is further evidenced by the presence of introgressions of these genes from *S. paradoxus* into *S. cerevisiae* in the Mediterranean oak lineage but not the European wine lineage (Almeida et al. 2017). The functional and ecological significance of these different alleles in vineyard versus oak forest environments remains an open question.

Taken together, the results of this study offer new insights into the target size, effect size, costs, and genetic identity of mutations conferring copper and sulfite resistance in *S. cerevisiae* and *S. paradoxus*. They also raise many important questions concerning the ecology of these species, and the explanatory and predictive utility of assaying the DME.

## Data availability statement

Raw reads produced in this study were deposited into NCBI under BioProject PRJNA1107929. The three colony size datasets used in this study (haploid, diploid, and sequenced) along with all of the accompanying metadata can be found at DOI: 10.6084/m9.figshare.25777512. This DOI also includes all images used for data collection of colony size and the processed data used to generate the figures.

## Acknowledgements

The authors would like to acknowledge Douda Bensasson for providing the natural isolates used to generate the focal strains used in this study. The authors would also like to acknowledge James Miller and other members of the Fay lab for helpful and thorough feedback on the writing of this work. Also, the authors acknowledge and thank Elaine Sia for allowing access to her microscope for the tetrad dissections performed in this study. Lastly, the authors would like to thank Nancy Chen, Elizabeth Grayhack, and Allen Orr for feedback and suggestions on this work.

## References

Adamo GM, Lotti M, Tamas MJ, Brocca S. 2012. Amplification of the *CUP1* gene is associated with evolution of copper tolerance in *Saccharomyces cerevisiae*. Microbiology. 158(9):2325–35.

Alexander RM. 1985. The ideal and the feasible: physical constraints on evolution. Biol J Linn Soc. 26(4):345–58.

Almeida P, Barbosa R, Bensasson D, Gonçalves P, Sampaio JP. 2017. Adaptive divergence in wine yeasts and their wild relatives suggests a prominent role for introgressions and rapid evolution at noncoding sites. Mol Ecol. 26(7):2167–82.

Almeida P, Barbosa R, Zalar P, Imanishi Y, Shimizu K, Turchetti B, Legras J, Serra M, Dequin S, Cou-loux A, et al. 2015. A population genomics insight into the Mediterranean origins of wine yeast do-mestication. Mol Ecol. 24(21):5412–27.

Amerine MA, Berg HW, Cruness WV. 1972. The technology of wine making 3rd edition. AVI Publishing Company, West Port. Connecticut.

Araya CL, Payen C, Dunham MJ, Fields S. 2010. Whole-genome sequencing of a laboratory-evolved yeast strain. BMC Genomics. 11:1–10.

Avery SV, Howlett NG, Radice S. 1996. Copper toxicity towards *Saccharomyces cerevisiae*: dependence on plasma membrane fatty acid composition. Appl Environ Microbiol. 62(11):3960–6.

Ayres PG. 2004. Alexis Millardet: France’s forgotten mycologist. Mycologist. 18(1):23–6.

Baryshnikova A, Costanzo M, Kim Y, Ding H, Koh J, Toufighi K, Youn J, Ou J, San Luis B, Bandyopadhyay S, et al. 2010. Quantitative analysis of fitness and genetic interactions in yeast on a genome scale. Nature Methods. 7(12):1017–24.

Besnard E, Chenu C, Robert M. 2001. Influence of organic amendments on copper distribution among particle-size and density fractions in Champagne vineyard soils. Environmental Pollution. 112(3):329–37.

Bharucha N, Ma J, Dobry CJ, Lawson SK, Yang Z, Kumar A. 2008. Analysis of the yeast kinome reveals a network of regulated protein localization during filamentous growth. Mol Biol Cell. 19(7):2708–17.

Birrell GW, Giaever G, Chu AM, Davis RW, Brown JM. 2001. A genome-wide screen in *Saccharomyces cerevisiae* for genes affecting UV radiation sensitivity. Proceedings of the National Academy of Sciences. 98(22):12608–13.

Blount ZD, Borland CZ, Lenski RE. 2008. Historical contingency and the evolution of a key innovation in an experimental population of *Escherichia coli*. Proceedings of the National Academy of Sciences. 105(23):7899–906.

Böndel KB, Kraemer SA, Samuels T, McClean D, Lachapelle J, Ness RW, Colegrave N, Keightley PD. 2019. Inferring the distribution of fitness effects of spontaneous mutations in *Chlamydomonas reinhardtii*. PLoS Biology. 17(6):e3000192.

Borchers HW. 2019. Package ‘pracma’. Practical Numerical Math Functions, Version. 2(5).

Borkow G, Gabbay J. 2009. Copper, an ancient remedy returning to fight microbial, fungal and viral infections. Current Chemical Biology. 3(3):272–8.

Bradshaw AD. 1991. The Croonian lecture, 1991. Genostasis and the limits to evolution. Philosophical Transactions of the Royal Society of London.Series B: Biological Sciences. 333(1267):289–305.

Bradshaw AD. 1984. The importance of evolutionary ideas in ecology and vice versa. Evol Ecol. :1–25.

Brewer BJ, Payen C, Raghuraman MK, Dunham MJ. 2011. Origin-dependent inverted-repeat amplification: A replication-based model for generating palindromic amplicons. PLoS Genetics. 7(3):e1002016.

Capdevila M, Bofill R, Palacios O, Atrian S. 2012. State-of-the-art of metallothioneins at the beginning of the 21st century. Coord Chem Rev. 256(1-2):46–62.

Carabelli AM, Peacock TP, Thorne LG, Harvey WT, Hughes J, COVID-19 Genomics UK Consortium, Peacock SJ, Barclay WS, de Silva TI, Towers GJ, et al. 2023. SARS-CoV-2 variant biology: Immune escape, transmission and fitness. Nature Reviews Microbiology. 21(3):162–77.

Caudy AA, Guan Y, Jia Y, Hansen C, DeSevo C, Hayes AP, Agee J, Alvarez-Dominguez JR, Arellano H, Barrett D, et al. 2013. A new system for comparative functional genomics of *Saccharomyces* yeasts. Genetics. 195(1):275–87.

Chang S, Lai H, Tung S, Leu J. 2013. Dynamic large-scale chromosomal rearrangements fuel rapid adaptation in yeast populations. PLoS Genetics. 9(1):e1003232.

Clark AC, Grant-Preece P, Cleghorn N, Scollary GR. 2015. Copper (II) addition to white wines containing hydrogen sulfide: residual copper concentration and activity. Australian Journal of Grape and Wine Research. 21(1):30–9.

Clowers KJ, Heilberger J, Piotrowski JS, Will JL, Gasch AP. 2015. Ecological and genetic barriers differentiate natural populations of *Saccharomyces cerevisiae*. Mol Biol Evol. 32(9):2317–27.

Couce A, Limdi A, Magnan M, Owen SV, Herren CM, Lenski RE, Tenaillon O, Baym M. 2024. Changing fitness effects of mutations through long-term bacterial evolution. Science. 383(6681):eadd1417.

Cowie RH, Bouchet P, Fontaine B. 2022. The sixth mass extinction: Fact, fiction or speculation? Biological Reviews. 97(2):640–63.

Culotta VC, Howard WR, Liu XF. 1994. *CRS5* encodes a metallothionein-like protein in *Saccharomyces cerevisiae*. J Biol Chem. 269(41):25295–302.

Cyert MS, Philpott CC. 2013. Regulation of cation balance in *Saccharomyces cerevisiae*. Genetics. 193(3):677–713.

Dashko S, Liu P, Volk H, Butinar L, Piškur J, Fay JC. 2016. Changes in the relative abundance of two *Saccharomyces* species from oak forests to wine fermentations. Frontiers in Microbiology. 7:215.

de Visser JAG, Rozen DE. 2006. Clonal interference and the periodic selection of new beneficial mutations in *Escherichia coli*. Genetics. 172(4):2093–100.

Dell’Amico E, Mazzocchi M, Cavalca L, Allievi L, Andreoni V. 2008. Assessment of bacterial community structure in a long-term copper-polluted ex-vineyard soil. Microbiol Res. 163(6):671–83.

Divol B, du Toit M, Duckitt E. 2012. Surviving in the presence of sulphur dioxide: strategies developed by wine yeasts. Appl Microbiol Biotechnol. 95:601–13.

Dixon B. 2004. Pushing Bordeaux mixture. The Lancet Infectious Diseases. 4(9):594.

Dobzhansky T, Ayala FJ, Stebbins GL, Valentine JW. 1977. Evolution. San Francisco: W. H. Freeman.

Dudley AM, Janse DM, Tanay A, Shamir R, Church GM. 2005. A global view of pleiotropy and phenotypically derived gene function in yeast. Molecular Systems Biology. 1(1):2005.0001.

Echave J, Barral M, Fraga-Corral M, Prieto MA, Simal-Gandara J. 2021. Bottle aging and storage of wines: A review. Molecules. 26(3):713.

Engle EK, Fay JC. 2012. Divergence of the yeast transcription factor *FZF1* affects sulfite resistance. PLoS Genetics. 8(6):e1002763.

Eyre-Walker A, Keightley PD. 2007. The distribution of fitness effects of new mutations. Nature Reviews Genetics. 8(8):610–8.

Falconer DS, Mackay TF. 1983. Quantitative genetics. Longman London.

Fay JC, McCullough HL, Sniegowski PD, Eisen MB. 2004. Population genetic variation in gene expression is associated with phenotypic variation in *Saccharomyces cerevisiae*. Genome Biol. 5:1–14.

Fernández-Calviño D, Bååth E. 2016. Interaction between pH and Cu toxicity on fungal and bacterial performance in soil. Soil Biol Biochem. 96:20-9. ffrench-Constant RH. 1994. The molecular and population genetics of cyclodiene insecticide resistance. Insect Biochem Mol Biol. 24(4):335–45.

Fisher RA. 1930. The genetical theory of natural selection. Oxford: Oxford University Press.

Fleiss A, O’Donnell S, Fournier T, Lu W, Agier N, Delmas S, Schacherer J, Fischer G. 2019. Reshuffling yeast chromosomes with CRISPR/Cas9. PLoS Genetics. 15(8):e1008332.

Fogel S, Welch JW. 1982. Tandem gene amplification mediates copper resistance in yeast. Proceedings of the National Academy of Sciences. 79(17):5342–6.

Fogel S, Welch JW, Cathala G, Karin M. 1983. Gene amplification in yeast: *CUP1* copy number regulates copper resistance. Curr Genet. 7:347–55.

Futuyma DJ. 2010. Evolutionary constraint and ecological consequences. Evolution. 64(7):1865–84.

Futuyma DJ. 1979. Evolutionary biology. Sunderland, MA: Sinauer.

Gagneux S, Long CD, Small PM, Van T, Schoolnik GK, Bohannan BJ. 2006. The competitive cost of antibiotic resistance in *Mycobacterium tuberculosis*. Science. 312(5782):1944–6.

García-Ríos E, Nuévalos M, Barrio E, Puig S, Guillamón JM. 2019. A new chromosomal rearrangement improves the adaptation of wine yeasts to sulfite. Environ Microbiol. 21(5):1771–81.

Gerrish PJ, Lenski RE. 1998. The fate of competing beneficial mutations in an asexual population. Genetica. 102:127–44.

Gerstein AC, Berman J. 2015. Shift and adapt: The costs and benefits of karyotype variations. Curr Opin Microbiol. 26:130–6.

Gerstein AC, Lo DS, Otto SP. 2012. Parallel genetic changes and nonparallel gene–environment interactions characterize the evolution of drug resistance in yeast. Genetics. 192(1):241–52.

Gerstein AC, Chun HE, Grant A, Otto SP. 2006. Genomic convergence toward diploidy in *Saccharomyces cerevisiae*. PLoS Genetics. 2(9):e145.

Gerstein AC, Ono J, Lo DS, Campbell ML, Kuzmin A, Otto SP. 2015. Too much of a good thing: The unique and repeated paths toward copper adaptation. Genetics. 199(2):555–71.

Gessler C, Pertot I, Perazzolli M. 2011. *Plasmopara viticola*: A review of knowledge on downy mildew of grapevine and effective disease management. Phytopathol Mediterr. 50(1):3–44.

Gietz RD, Schiestl RH. 2007. High-efficiency yeast transformation using the LiAc/SS carrier DNA/PEG method. Nature Protocols. 2(1):31–4.

Gilchrist C, Stelkens R. 2019. Aneuploidy in yeast: Segregation error or adaptation mechanism? Yeast. 36(9):525–39.

Gillespie JH. 1991. The causes of molecular evolution. Oxford: Oxford University Press.

Goldstein AL, McCusker JH. 1999. Three new dominant drug resistance cassettes for gene disruption in *Saccharomyces cerevisiae*. Yeast. 15(14):1541–53.

Gonzalez A, Bell G. 2013. Evolutionary rescue and adaptation to abrupt environmental change depends upon the history of stress. Philosophical Transactions of the Royal Society B: Biological Sciences. 368(1610):20120079.

Good BH, Desai MM. 2015. The impact of macroscopic epistasis on long-term evolutionary dynamics. Genetics. 199(1):177–90.

Goto-Yamamoto N, Kitano K, Shiki K, Yoshida Y, Suzuki T, Iwata T, Yamane Y, Hara S. 1998. *SSU1R*, a sulfite resistance gene of wine yeast, is an allele of *SSU1* with a different upstream sequence. J Ferment Bioeng. 86(5):427–33.

Grangeteau C, David V, Herve A, Guilloux-Benatier M, Rousseaux S. 2017. The sensitivity of yeasts and yeasts-like fungi to copper and sulfur could explain lower yeast biodiversity in organic vineyards. FEMS Yeast Research. 17(8):fox092.

Gruber JD, Vogel K, Kalay G, Wittkopp PJ. 2012. Contrasting properties of gene-specific regulatory, coding, and copy number mutations in *Saccharomyces cerevisiae*: frequency, effects, and dominance. PLoS Genetics. 8(2):e1002497.

Guarini N, Saliba E, André B. 2024. Phosphoregulation of the yeast Pma1 H -ATPase autoinhibitory domain involves the Ptk1/2 kinases and the Glc7 PP1 phosphatase and is under TORC1 control. Plos Genetics. 20(1):e1011121.

Hill WG, Caballero A. 1992. Artificial selection experiments. Annu Rev Ecol Syst. 23(1):287–310.

Hodgins-Davis A, Duveau F, Walker EA, Wittkopp PJ. 2019. Empirical measures of mutational effects define neutral models of regulatory evolution in *Saccharomyces cerevisiae*. Proceedings of the National Academy of Sciences. 116(42):21085–93.

Hoffmann AA, Hallas RJ, Dean JA, Schiffer M. 2003. Low potential for climatic stress adaptation in a rainforest *Drosophila* species. Science. 301(5629):100–2.

Hull RM, Cruz C, Jack CV, Houseley J. 2017. Environmental change drives accelerated adaptation through stimulated copy number variation. PLoS Biology. 15(6):e2001333.

Huxley C, Green ED, Dunham I. 1990. Rapid assessment of *S. cerevisiae* mating type by PCR. Trends in Genetics: TIG. 6(8):236.

Hyma KE, Fay JC. 2013. Mixing of vineyard and oak-tree ecotypes of *Saccharomyces cerevisiae* in North American vineyards. Mol Ecol. 22(11):2917–30.

Jolly NP, Augustyn O, Pretorius IS. 2006. The role and use of non-*Saccharomyces* yeasts in wine production. South African Journal of Enology & Viticulture. .

Kapitzky L, Beltrao P, Berens TJ, Gassner N, Zhou C, Wüster A, Wu J, Babu MM, Elledge SJ, Toczyski D, et al. 2010. Cross-species chemogenomic profiling reveals evolutionarily conserved drug mode of action. Molecular Systems Biology. 6(1):451.

Karasov T, Messer PW, Petrov DA. 2010. Evidence that adaptation in *Drosophila* is not limited by mutation at single sites. PLoS Genetics. 6(6):e1000924.

Komárek M, Száková J, Rohošková M, Javorská H, Chrastný V, Balík J. 2008. Copper contamination of vineyard soils from small wine producers: A case study from the Czech Republic. Geoderma. 147(1-2):16–22.

Ladrey C. 1871. L’art de faire le vin. Dijon, France: Imprimerie J.E. Rabutot.

Lang GI, Murray AW. 2008. Estimating the per-base-pair mutation rate in the yeast *Saccharomyces cerevisiae*. Genetics. 178(1):67–82.

Lewontin RC. 1974. The genetic basis of evolutionary change. Columbia University Press New York.

Li H. 2013. Aligning sequence reads, clone sequences and assembly contigs with BWA-MEM. arXiv Preprint arXiv:1303.3997. .

Linderholm AL, Findleton CL, Kumar G, Hong Y, Bisson LF. 2008. Identification of genes affecting hydrogen sulfide formation in *Saccharomyces cerevisiae*. Appl Environ Microbiol. 74(5):1418–27.

Liti G, Carter DM, Moses AM, Warringer J, Parts L, James SA, Davey RP, Roberts IN, Burt A, Koufopanou V, et al. 2009. Population genomics of domestic and wild yeasts. Nature. 458(7236):337–41.

Mable BK, Otto SP. 2001. Masking and purging mutations following EMS treatment in haploid, diploid and tetraploid yeast (*Saccharomyces cerevisiae*). Genetics Research. 77(1):9–26.

Macías LG, Flores MG, Adam AC, Rodríguez ME, Querol A, Barrio E, Lopes CA, Pérez-Torrado R. 2021. Convergent adaptation of *Saccharomyces uvarum* to sulfite, an antimicrobial preservative widely used in human-driven fermentations. PLoS Genetics. 17(11):e1009872.

Mackie KA, Müller T, Zikeli S, Kandeler E. 2013. Long-term copper application in an organic vineyard modifies spatial distribution of soil micro-organisms. Soil Biol Biochem. 65:245–53.

Mackie KA, Müller T, Kandeler E. 2012. Remediation of copper in vineyards–a mini review. Environmental Pollution. 167:16–26.

Macomber L, Imlay JA. 2009. The iron-sulfur clusters of dehydratases are primary intracellular targets of copper toxicity. Proceedings of the National Academy of Sciences. 106(20):8344–9.

Maddamsetti R, Lenski RE, Barrick JE. 2015. Adaptation, clonal interference, and frequency-dependent interactions in a long-term evolution experiment with *Escherichia coli*. Genetics. 200(2):619–31.

Mao P, Wyrick JJ, Roberts SA, Smerdon MJ. 2017. UV-induced DNA damage and mutagenesis in chromatin. Photochem Photobiol. 93(1):216–28.

Masson E. 1887. La bouille Bourguignonne. Progess Agricole Et Viticole. 1:511–3.

Mazón MJ, Eraso P, Portillo F. 2015. Specific phosphoantibodies reveal two phosphorylation sites in yeast Pma1 in response to glucose. FEMS Yeast Research. 15(5):fov030.

Metzger BP, Duveau F, Yuan DC, Tryban S, Yang B, Wittkopp PJ. 2016. Contrasting frequencies and effects of cis-and trans-regulatory mutations affecting gene expression. Mol Biol Evol. 33(5):1131–46.

Millardet A. 1885. Traitement du mildiou par le mélange de sulphate de cuivre et chaux. Journal Agriculture Pratique. 49(2):707–10.

Miller JH, Fasanello VJ, Liu P, Longan ER, Botero CA, Fay JC. 2022. Using colony size to measure fitness in *Saccharomyces cerevisiae*. PLoS One. 17(10):e0271709.

Money NP. 2006. The triumph of the fungi: A rotten history. Oxford University Press.

Morsomme P, Slayman CW, Goffeau A. 2000. Mutagenic study of the structure, function and biogenesis of the yeast plasma membrane H -ATPase. Biochimica Et Biophysica Acta (BBA)-Reviews on Biomembranes. 1469(3):133–57.

Mortimer RK, Johnston JR. 1986. Genealogy of principal strains of the yeast genetic stock center. Genetics. 113(1):35–43.

Onetto CA, Kutyna DR, Kolouchova R, McCarthy J, Borneman AR, Schmidt SA. 2023. SO_2_ and copper tolerance exhibit an evolutionary trade-off in *Saccharomyces cerevisiae*. PLoS Genetics. 19(3):e1010692.

Orr HA. 2005. The genetic theory of adaptation: A brief history. Nature Reviews Genetics. 6(2):119–27.

Orr HA. 2003. The distribution of fitness effects among beneficial mutations. Genetics. 163(4):1519–26.

Orr HA. 1998. The population genetics of adaptation: The distribution of factors fixed during adaptive evolution. Evolution. 52(4):935–49.

Orr HA, Unckless RL. 2014. The population genetics of evolutionary rescue. PLoS Genetics. 10(8):e1004551.

Park H, Lopez NI, Bakalinsky AT. 1999. Use of sulfite resistance in *Saccharomyces cerevisiae* as a dominant selectable marker. Curr Genet. 36:339–44.

Pecci A, Borgna E, Mileto S, Dalla Longa E, Bosi G, Florenzano A, Mercuri AM, Corazza S, Marchesini M, Vidale M. 2020. Wine consumption in bronze age Italy: Combining organic residue analysis, botanical data and ceramic variability. Journal of Archaeological Science. 123:105256.

Pentz JT, Lind PA. 2021. Forecasting of phenotypic and genetic outcomes of experimental evolution in *Pseudomonas protegens*. PLoS Genetics. 17(8):e1009722.

Pentz JT, Biswas A, Alsaed B, Lind PA. 2024. Forecasting of phenotypic and genetic outcomes of experimental evolution in *Pseudomonas syringae* and *Pseudomonas savastanoi*. bioRxiv. :2024.02. 10.579745.

Pérez-Ortın JE, Querol A, Puig S, Barrio E. 2002. Molecular characterization of a chromosomal rearrangement involved in the adaptive evolution of yeast strains. Genome Res. 12(10):1533–9.

Perfeito L, Fernandes L, Mota C, Gordo I. 2007. Adaptive mutations in bacteria: high rate and small effects. Science. 317(5839):813–5.

Redžepović S, Orlić S, Sikora S, Majdak A, Pretorius IS. 2002. Identification and characterization of *Saccharomyces cerevisiae* and *Saccharomyces paradoxus* strains isolated from Croatian vineyards. Lett Appl Microbiol. 35(4):305–10.

Rey O, Danchin E, Mirouze M, Loot C, Blanchet S. 2016. Adaptation to global change: A transposable element–epigenetics perspective. Trends in Ecology & Evolution. 31(7):514–26.

Robinson HA, Pinharanda A, Bensasson D. 2016. Summer temperature can predict the distribution of wild yeast populations. Ecology and Evolution. 6(4):1236–50.

Sanchez MR, Miller AW, Liachko I, Sunshine AB, Lynch B, Huang M, Alcantara E, DeSevo CG, Pai DA, Tucker CM, et al. 2017. Differential paralog divergence modulates genome evolution across yeast species. PLoS Genetics. 13(2):e1006585.

Sanjuán R, Moya A, Elena SF. 2004. The distribution of fitness effects caused by single-nucleotide substitutions in an RNA virus. Proceedings of the National Academy of Sciences. 101(22):8396–401.

Scannell DR, Zill OA, Rokas A, Payen C, Dunham MJ, Eisen MB, Rine J, Johnston M, Hittinger CT. 2011. The awesome power of yeast evolutionary genetics: New genome sequences and strain resources for the *Saccharomyces sensu stricto* genus. G3: Genes| Genomes| Genetics. 1(1):11–25.

Selmecki AM, Maruvka YE, Richmond PA, Guillet M, Shoresh N, Sorenson AL, De S, Kishony R, Michor F, Dowell R, et al. 2015. Polyploidy can drive rapid adaptation in yeast. Nature. 519(7543):349–52.

Shu Y, Yang H, Hallberg E, Hallberg R. 1997. Molecular genetic analysis of Rts1p, a B′ regulatory subunit of *Saccharomyces cerevisiae* protein phosphatase 2A. Mol Cell Biol. .

Sniegowski PD, Dombrowski PG, Fingerman E. 2002. *Saccharomyces cerevisiae* and *Saccharomyces paradoxus* coexist in a natural woodland site in North America and display different levels of reproductive isolation from European conspecifics. FEMS Yeast Research. 1(4):299–306.

Stratford M, Morgan P, Rose AH. 1987. Sulphur dioxide resistance in *Saccharomyces cerevisiae* and *Saccharomycodes ludwigii*. J Gen Microbiol. 133(8):2173–9.

Strope PK, Skelly DA, Kozmin SG, Mahadevan G, Stone EA, Magwene PM, Dietrich FS, McCusker JH. 2015. The 100-genomes strains, an *S. cerevisiae* resource that illuminates its natural phenotypic and genotypic variation and emergence as an opportunistic pathogen. Genome Res. 25(5):762–74.

Tkeshelashvili LK, McBride T, Spence K, Loeb LA. 1991. Mutation spectrum of copper-induced DNA damage. J Biol Chem. 266(10):6401–6.

Todd RT, Selmecki A. 2020. Expandable and reversible copy number amplification drives rapid adaptation to antifungal drugs. Elife. 9:e58349.

Umekawa M, Klionsky DJ. 2012. Ksp1 kinase regulates autophagy via the target of rapamycin complex 1 (TORC1) pathway. J Biol Chem. 287(20):16300–10.

Valero E, Tronchoni J, Morales P, Gonzalez R. 2020. Autophagy is required for sulfur dioxide tolerance in *Saccharomyces cerevisiae*. Microbial Biotechnology. 13(2):599–604.

van Bakel H, Strengman E, Wijmenga C, Holstege FC. 2005. Gene expression profiling and phenotype analyses of *S. cerevisiae* in response to changing copper reveals six genes with new roles in copper and iron metabolism. Physiological Genomics. 22(3):356–67.

Van den Bergh B, Swings T, Fauvart M, Michiels J. 2018. Experimental design, population dynamics, and diversity in microbial experimental evolution. Microbiology and Molecular Biology Reviews. 82(3):10.1128/mmbr.00008-18.

Varela C, Bartel C, Roach M, Borneman A, Curtin C. 2019. *Brettanomyces bruxellensis SSU1* haplotypes confer different levels of sulfite tolerance when expressed in a *Saccharomyces cerevisiae SSU1* null mutant. Appl Environ Microbiol. 85(4):2429.

Vaudano E, Quinterno G, Costantini A, Pulcini L, Pessione E, Garcia-Moruno E. 2019. Yeast distribution in Grignolino grapes growing in a new vineyard in Piedmont and the technological characterization of indigenous *Saccharomyces* spp. strains. Int J Food Microbiol. 289:154–61.

Vela E, Hernández-Orte P, Franco-Luesma E, Ferreira V. 2017. The effects of copper fining on the wine content in sulfur off-odors and on their evolution during accelerated anoxic storage. Food Chem. 231:212–21.

Vogwill T, Kojadinovic M, Maclean RC. 2016. Epistasis between antibiotic resistance mutations and genetic background shape the fitness effect of resistance across species of *Pseudomonas*. Proceedings of the Royal Society B: Biological Sciences. 283(1830):20160151.

Vogwill T, Kojadinovic M, Furió V, MacLean RC. 2014. Testing the role of genetic background in parallel evolution using the comparative experimental evolution of antibiotic resistance. Mol Biol Evol. 31(12):3314–23.

Warringer J, Zörgö E, Cubillos FA, Zia A, Gjuvsland A, Simpson JT, Forsmark A, Durbin R, Omholt SW, Louis EJ, et al. 2011. Trait variation in yeast is defined by population history. PLoS Genetics. 7(6):e1002111.

Welch J, Fogel S, Buchman C, Karin M. 1989. The *CUP2* gene product regulates the expression of the *CUP1* gene, coding for yeast metallothionein. Embo J. 8(1):255–60.

Welch JW, Fogel S, Cathala G, Karin M. 1983. Industrial yeasts display tandem gene iteration at the *CUP1* region. Mol Cell Biol. 3(8):1353–61.

Williams KM, Liu P, Fay JC. 2015. Evolution of ecological dominance of yeast species in high-sugar environments. Evolution. 69(8):2079–93.

Williams-Hart T, Wu X, Tatchell K. 2002. Protein phosphatase type 1 regulates ion homeostasis in *Saccharomyces cerevisiae*. Genetics. 160(4):1423–37.

Wloch DM, Szafraniec K, Borts RH, Korona R. 2001. Direct estimate of the mutation rate and the distribution of fitness effects in the yeast *Saccharomyces cerevisiae*. Genetics. 159(2):441–52.

Wu NC, Wilson IA. 2017. A perspective on the structural and functional constraints for immune evasion: insights from influenza virus. J Mol Biol. 429(17):2694–709.

Yona AH, Manor YS, Herbst RH, Romano GH, Mitchell A, Kupiec M, Pilpel Y, Dahan O. 2012. Chromosomal duplication is a transient evolutionary solution to stress. Proceedings of the National Academy of Sciences. 109(51):21010–5.

Yoshikawa K, Tanaka T, Furusawa C, Nagahisa K, Hirasawa T, Shimizu H. 2009. Comprehensive phenotypic analysis for identification of genes affecting growth under ethanol stress in *Saccharomyces cerevisiae*. FEMS Yeast Research. 9(1):32–44.

Young BP, Shin JJ, Orij R, Chao JT, Li SC, Guan XL, Khong A, Jan E, Wenk MR, Prinz WA, et al. 2010. Phosphatidic acid is a pH biosensor that links membrane biogenesis to metabolism. Science. 329(5995):1085–8.

Young M, Heit S, Bublitz M. 2023. Structure, function and biogenesis of the fungal proton pump Pma1. Biochimica Et Biophysica Acta (BBA)-Molecular Cell Research. :119600.

Yue J, Li J, Aigrain L, Hallin J, Persson K, Oliver K, Bergström A, Coupland P, Warringer J, Lagomarsino MC, et al. 2017. Contrasting evolutionary genome dynamics between domesticated and wild yeasts. Nat Genet. 49(6):913–24.

Zackrisson M, Hallin J, Ottosson L, Dahl P, Fernandez-Parada E, Ländström E, Fernandez-Ricaud L, Kaferle P, Skyman A, Stenberg S, et al. 2016. Scan-o-matic: high-resolution microbial phenomics at a massive scale. G3: Genes, Genomes, Genetics. 6(9):3003–14.

Zeyl C, Vanderford T, Carter M. 2003. An evolutionary advantage of haploidy in large yeast populations. Science. 299(5606):555–8.

Zhao Y, Dominska M, Petrova A, Bagshaw H, Kokoska RJ, Petes TD. 2017. Properties of mitotic and meiotic recombination in the tandemly-repeated *CUP1* gene cluster in the yeast *Saccharomyces cerevisiae*. Genetics. 206(2):785–800.

Zhao Y, Strope PK, Kozmin SG, McCusker JH, Dietrich FS, Kokoska RJ, Petes TD. 2014. Structures of naturally evolved *CUP1* tandem arrays in yeast indicate that these arrays are generated by unequal nonhomologous recombination. G3: Genes, Genomes, Genetics. 4(11):2259–69.

Zimmer A, Durand C, Loira N, Durrens P, Sherman DJ, Marullo P. 2014. QTL dissection of lag phase in wine fermentation reveals a new translocation responsible for *Saccharomyces cerevisiae* adaptation to sulfite. Plos One. 9(1):e86298.

